# Single fluorogen imaging reveals distinct environmental and structural features of biomolecular condensates

**DOI:** 10.1101/2023.01.26.525727

**Authors:** Tingting Wu, Matthew R. King, Yuanxin Qiu, Mina Farag, Rohit V. Pappu, Matthew D. Lew

## Abstract

Biomolecular condensates are viscoelastic materials. Simulations predict that fluid-like condensations are defined by spatially inhomogeneous organization of the underlying molecules. Here, we test these predictions using single-fluorogen tracking and super-resolution imaging. Specifically, we leverage the localization and orientational preferences of freely diffusing fluorogens and the solvatochromic effect whereby specific fluorogens are turned on in response to condensate microenvironments. We deployed three different fluorogens to probe the microenvironments and molecular organization of different protein-based condensates. The spatiotemporal resolution and environmental sensitivity afforded by single-fluorogen imaging shows that the internal environments of condensates are more hydrophobic than coexisting dilute phases. Molecules within condensates are organized in a spatially inhomogeneous manner, and this gives rise to slow-moving nanoscale molecular clusters that coexist with fast-moving molecules. Fluorogens that localize preferentially to the interface help us map their distinct features. Our findings provide a structural and dynamical basis for the viscoelasticity of condensates.

## Main Text

Biomolecular condensates are thought to provide spatial and temporal control over cellular matter ^1^. Proteins with intrinsically disordered regions (IDRs) such as prion-like low complexity domains (PLCDs) and arginine-glycine-rich (RG-rich) regions are prominent drivers of different types of condensates ^2, 3^. Several investigations have suggested that simple, single-component condensates formed by PLCDs and RG-rich IDRs and complex multicomponent condensates such as nucleoli and the mitochondrial nucleoid^4^ are viscoelastic materials ^5, 6, 7, 8, 9, 10^. Viscoelastic materials should feature network-like internal structures ^11, 12^. The dynamical rearrangement of these networks and the transport properties of the underlying molecules should determine whether the long-time behaviors of viscoelastic materials are dominantly viscous (fluid-like) or elastic (solid-like) ^10^. Despite the central importance of nano- and mesoscale organization of molecules to condensate viscoelasticity, these details remain elusive even within simple one-component condensates.

Recent computational studies have predicted that within simple, one-component condensates formed by PLCDs, the protein molecules organize to form spatially inhomogeneous networks ^13, 14^ (**Fig. 1a**). Results from simulations suggest that internal networks in condensates have dynamic, hub-and-spoke organization with hubs being nanoscale clusters of molecules that are generated by reversible physical crosslinks among molecules ^13, 14^. Here, we report results from experimental tests of computational predictions using single-molecule tracking and single-molecule orientation localization microscopy (SMOLM). In these experiments, we combine freely diffusing fluorogenic probes ^15, 16, 17^ with single-molecule imaging ^18, 19, 20, 21, 22, 23^ to characterize the internal solvent properties as well as the organization and dynamics of molecules within condensates.

**Fig. 1.**
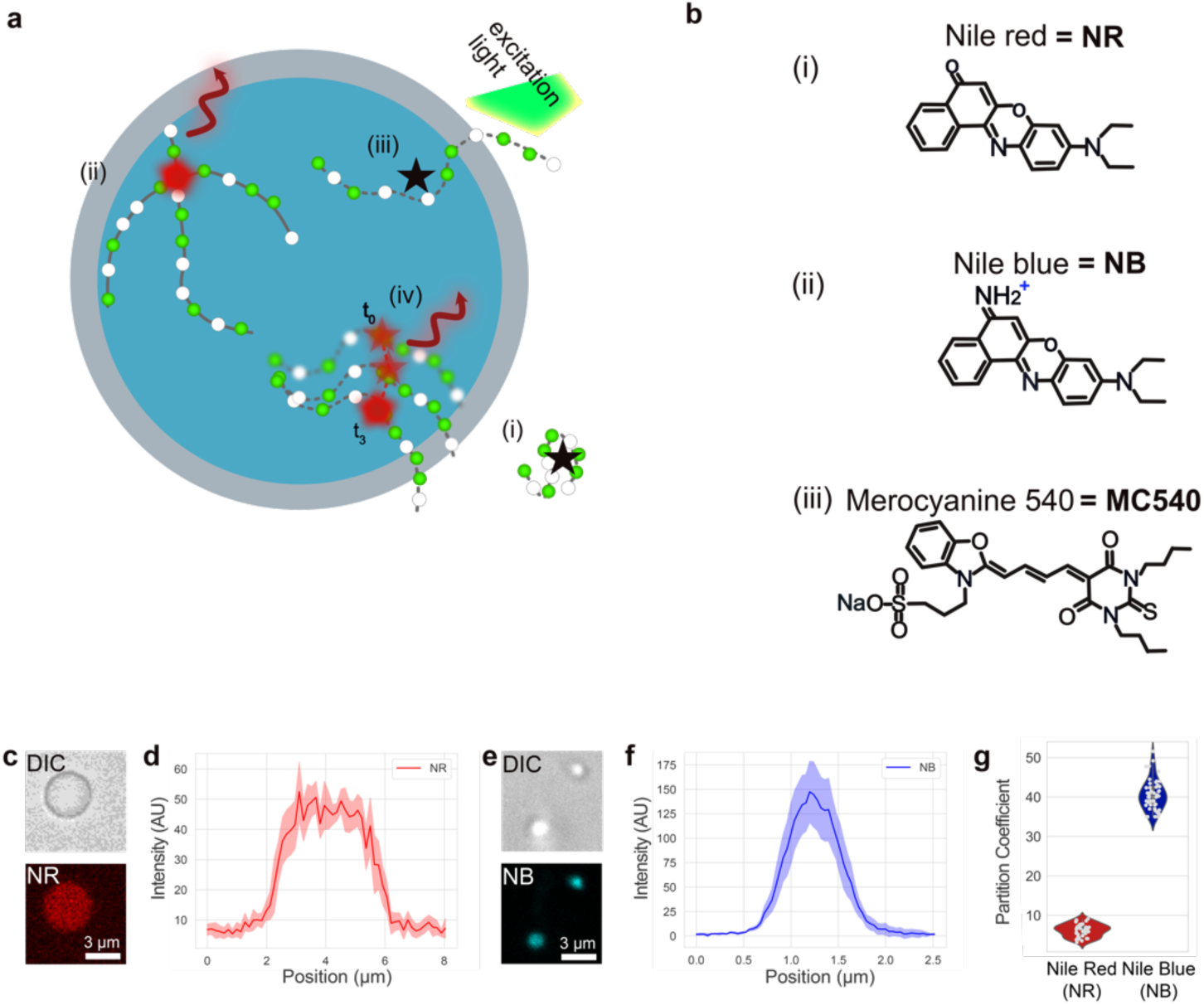
Details of fluorogenic probes and their partitioning into biomolecular condensates. **a**: As fluorogens (shown as black stars) diffuse through solution, they encounter different chemical environments in the dilute phase (white space), in the condensate (blue circle), and in the interface between the dilute phase and condensate (gray region). In response, they either (i, iii) remain dark (black stars) or (ii, iv) become fluorescent (red stars and curved arrows). Their (ii, iv) speeds of movement and brightness are also affected. **b**: Chemical structures of the three different fluorogens used for imaging condensates: (i) Nile red (NR), (ii) Nile blue (NB), and (iii) merocyanine 540 (MC540). (c) Condensates formed by unlabeled A1-LCD molecules were imaged using differential interference contrast (DIC) microscopy. NR partitions into these condensates (signal shown in the red channel). (d) Point scanning confocal microscopy was used to quantify the partition coefficients of NR into A1-LCD condensates. (e) DIC images of A1-LCD condensates and confocal images showing the partitioning of NB into these condensates. (f) Point scanning confocal microscopy was used to quantify the partition coefficients of NB into A1-LCD condensates. (g) Partition coefficients for NR and NB measured using partitioning data gathered from ∼50 different condensates. The median values are 6.5 and 40 for NR and NB, respectively.

Environmentally sensitive dyes change their fluorescence intensity (making them fluorogens) or color (making them solvatochromic) based on their microenvironment (**Extended Data** Fig. 1) ^15, 16, 17, 24, 25, 26, 27^. We deployed fluorogenic molecules that are solvatochromic and used them as structural probes in super-resolution single-molecule imaging to investigate condensates formed by three sequence variants of the PLCD of hnRNP-A1 (A1-LCD) ^2, 28^ and the RG-rich IDR from the protein DDX4 ^3, 29^. We focused on condensates formed by single proteins to test the structural basis for measured viscoelastic properties ^7, 30, 31^ and predictions that have emerged from physical theory and computations ^13, 14, 32^.

In separate measurements we used different polarity-sensing fluorogens namely, Nile blue (NB), Nile red (NR) ^15, 16^, and merocyanine 540 (MC540)^17^. These fluorogens are dark in aqueous solutions, even when they are pumped with an excitation laser. As the fluorogens diffuse through solution, they can either partition into hydrophobic environments or bind specifically to hydrophobic pockets within or outside condensates. This binding causes an increase in their fluorescence quantum yield and a shift in their absorption and emission spectra, termed solvatochromism (**Fig. 1a**).

NR (**Fig. 1b, (i)**) is minimally soluble in aqueous buffers, and it has a solubility of 280 µM (0.09 mg/ml) in a 1:10 mixture of DMSO (dimethyl sulfoxide) and phosphate buffered saline (PBS). In contrast, NB (**Fig. 1b, (ii)**), which is essentially the same as NR, albeit with an amine, has a solubility in water of 0.14 M (50 g/l). MC540 (**Fig. 1b, (iii)**) partitions to interfaces between membranes and aqueous environments ^17^. Interiors of condensates are likely to be microenvironments that are distinct from the coexisting phases ^4, 8, 13, 14, 33, 34^ that are delineated by interfaces. Therefore, we used MC540 as a probe of condensate interfaces. Note that none of the proteins are chemically modified by dyes or other tags. Instead, the fluorogenic dyes, which are present at low micromolar or sub-micromolar concentrations in single-molecule measurements, freely diffuse in solution and through the condensates ^35^.

Prior to deploying the fluorogens as imaging probes within condensate interiors, we used point scanning confocal microscopy to measure the partitioning of NR and NB into condensates formed by wild-type A1-LCD (**Fig. 1c-1f**). The partitioning measurements were performed using 34 µM NR and 40 µM NB. Across ∼50 distinct measurements we obtain median values of 6.5 and 40 for the partition coefficients of NR and NB into A1-LCD-rich phases (**Fig. 1g**). The partition coefficient of NB is eight times higher than that of NR. In the single-molecule measurements, we use concentrations of 3.4 µM for NR and 2 nM for NB. Based on the measured partition coefficients, the inferred concentration for NR and NB in the dense phase would be 17 µM and 0.08 µM, respectively. Therefore, the concentrations of NR and NB in the dense phase are well below the threshold concentrations that define their solubility limits phase separation in aqueous solvents.

We also asked if the fluorogens function as ligands that alter the driving forces for phase separation of the macromolecules. Ligands that stabilize dense phases will lower the macromolecular saturation concentration (*c*_sat_) whereas ligands that destabilize dense phases will have the opposite effect ^36, 37^. We measured the effects of fluorogens on the *c*_sat_ values of A1-LCD molecules. These measurements were performed at two different temperatures and different bulk concentrations of each of the dyes. The data show that effects of the dyes on the *c*_sat_ are negligible, being within the narrow errors of the measured *c*_sat_ values in the absence of the dyes (**Extended Data** Fig. 2a-b). Therefore, the fluorogenic and solvatochromic dyes are non-perturbative when used at single-molecule imaging concentrations. The control measurements establish that the features we extract from single-molecule measurements are direct consequences of interactions of the dyes with the macromolecules and not a consequence of the interactions of the dyes with themselves.

### Single-molecule imaging shows that spatial inhomogeneities extend down to the nanoscale

We leveraged the diffusion and transient binding of NB and NR ^35, 38^ as a photo-switching (“blinking”) mechanism for super-resolution, single-molecule localization microscopy (SMLM) ^39, 40, 41, 42^. As individual probes diffuse through solution, they remain dark until they encounter hydrophobic environments or bind to hydrophobic sites. Within hydrophobic pockets, they become bright, and if they are sufficiently immobilized within the pocket (e.g., ∼10 ms—the timescale of the camera integration time), then enough fluorescence signal can be accumulated to detect and localize single fluorogens. The fluorogens become dark when they leave hydrophobic pockets ^18^. Accumulating the positions of many blinking events yields detailed SMLM maps of internal structures within condensates with nanoscale resolution and single-molecule sensitivity (**Movie S1**).

Super-resolution images of single condensates show that both NR and NB exhibit non-uniform localization patterns (**Fig. 2a, 2b**). We measured digital on / off blinking of the fluorogens within apparent “hotspots” that are formed by the macromolecules within condensates. The hotspots are repeated “bursts” of single-molecule fluorescence turning on and off via binding and unbinding / photobleaching. Based on the observations in computations ^13^, we refer to the hotpots as hubs. We observed hubs at the nanoscale in SMLM images collected using both NR and NB (**Fig. 2c**). To quantify the spatial inhomogeneity using single-molecule blinking (**Fig. 2d**), we calculated the excess variance of the number of single-fluorogen localizations within a spatial pixel within each condensate (see Methods). If a fluorogenic probe has a uniform probability of blinking across the entire condensate, e.g., in the absence of any hotspots, then its localization density would be Poisson-distributed ^43^. This behavior yields an excess variance of zero (**Fig. S3**). The condensates imaged sequentially by NB and NR showed larger than expected variances with a mean excess variance of 1.3 ± 0.8 (std. dev., 2.1×10^4^ localizations/condensate on average). Thus, SMLM shows that both NB and NR exhibit large and significant spatial variations in their blinking dynamics (**Fig. 2d**).

**Fig. 2.**
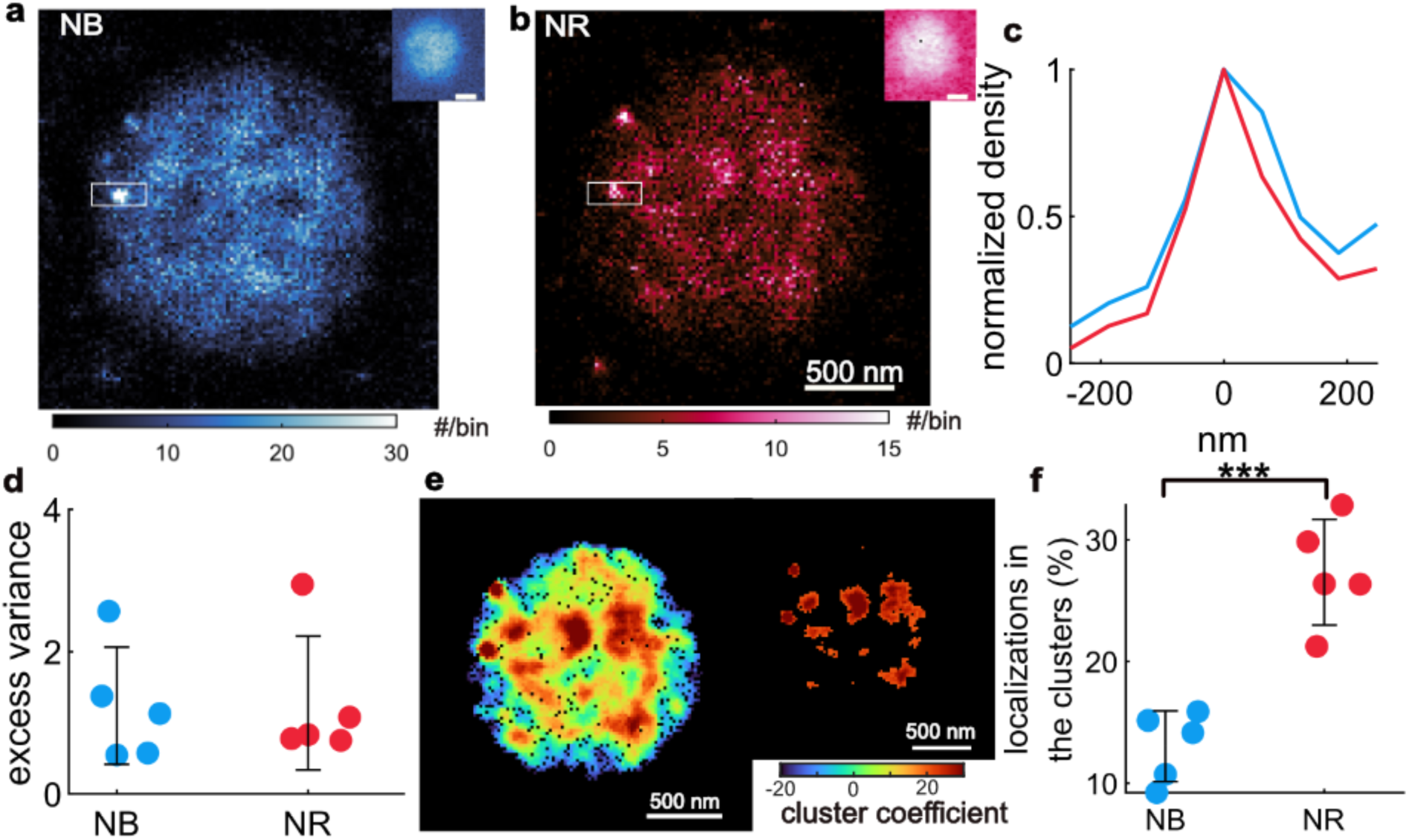
Single molecule localization microscopy (SMLM) of NB and NR reveals that they bind to molecules that are organized into nanoscale clusters within condensates. **a**, **b**: SMLM images of a single A1-LCD condensate collected using (**a**) NB and (**b**) NR. Top insets: epifluorescence images. Color bars: number (#) of single-molecule localizations within each 20 nm × 20 nm bin. **c**: Localization profile along the long axis of the white box shown in (**a, b**). **d**: Quantifying the binding and activation of NB (blue) and NR (red) within five condensates using excess variance. Larger excess variance values represent greater heterogeneities in the blinking statistics of fluorogenic probes within a condensate; zero excess variance represents uniform blinking statistics throughout the condensate. **e**: A1-LCD condensate in (**b**) imaged by NR and color-coded by clustering coefficient; molecules with clustering coefficients above a threshold of 20 are classified as being clustered. Inset: map of regions that contain clustered NR localizations. **f**: Percentages of single-molecule localizations that are spatially clustered for five condensates.

The super-resolution images of NR show the presence of nanoscale hubs (**Fig. 2c**). Next, we measured the relative proportion of localizations that are spatially clustered to quantify the relative affinities of NR and NB for localizing to nanoscale hubs (**Fig. 2e, 2f)**. Our analysis is motivated by theories showing that density fluctuations are useful order parameters to assay for density inhomogeneities ^44^. For each localization, we compared the local and surrounding density and then computed a clustering coefficient ^45, 46^. Molecules with clustering coefficients above a threshold of 20 are classified as being clustered (**Fig. 2e**, see Methods). For the five condensates imaged by NR and NB, 27% ± 3% of NR molecules and 13% ± 4% of NB molecules were clustered. This indicates that clustering is detected twice as often with NR when compared to NB. The implication is that while both probes exhibit a strong affinity to hubs, NR localizes more strongly to nanoscale clusters than NB.

Next, we quantified the similarities and differences between spatial inhomogeneities mapped by NR and NB ^47, 48^. For images collected using both NR and NB, we found that the diffuse regions, characterized by small relative single-fluorogen localization densities, are mostly uncorrelated, i.e., random. However, regions exhibiting high NR and NB blinking events are highly correlated with one another (**Extended Data** Fig. 4). These data suggest that the affinities and binding lifetimes of the fluorogens at hubs are dye-specific, but the hubs identified by the dyes are equivalent.

### Single-molecule tracking shows slow moving nanoscale hubs that coexist with fast moving molecules

Our data indicate that nanoscale hubs represent clusters of proteins that define distinctive local environments. Single-molecule imaging also shows that nanoscale hubs within each of the condensates disappear from one location and reappear at nearby locations (**Movies S2 and S3**). We probed these dynamics inside condensates by tracking the movements and fluorescence burst durations of individual fluorogens (**Movie S4**). The burst duration *t*_b_ is correlated with the amount of time a fluorogen spends in a fluorescence-promoting environment. The speed measures the distance covered by a fluorogen within 10 ms intervals (**Fig. 3a**, **Extended Data** Fig. 5). As a fluorogen explores a condensate, changes in the local environment cause both its burst duration and speed of movement to vary. For example, the burst durations of NR range from 10 ms to more than 170 ms (**Fig. 3b**). This heterogeneity is highlighted by reconstructing SMLM images using only single molecules with short burst durations (*t*_b_ ≤ 60 ms, **Fig. 3c**) versus those with long burst durations (*t*_b_ > 60 ms, **Fig. 3d**). Single molecules with short burst durations are distributed uniformly across the condensate, while hotspots are dominant in the long-burst duration images. This trend was also observed when the condensates were imaged using NB (**Extended Data** Fig. 5i-5k). However, when compared to NR, NB exhibits larger displacements (**Extended Data** Fig. 5b,5d). These data are consistent with the observation that the NB dyes localize to equivalent hubs within condensates but their dwell times at the hubs are different from those of NR.

**Fig. 3.**
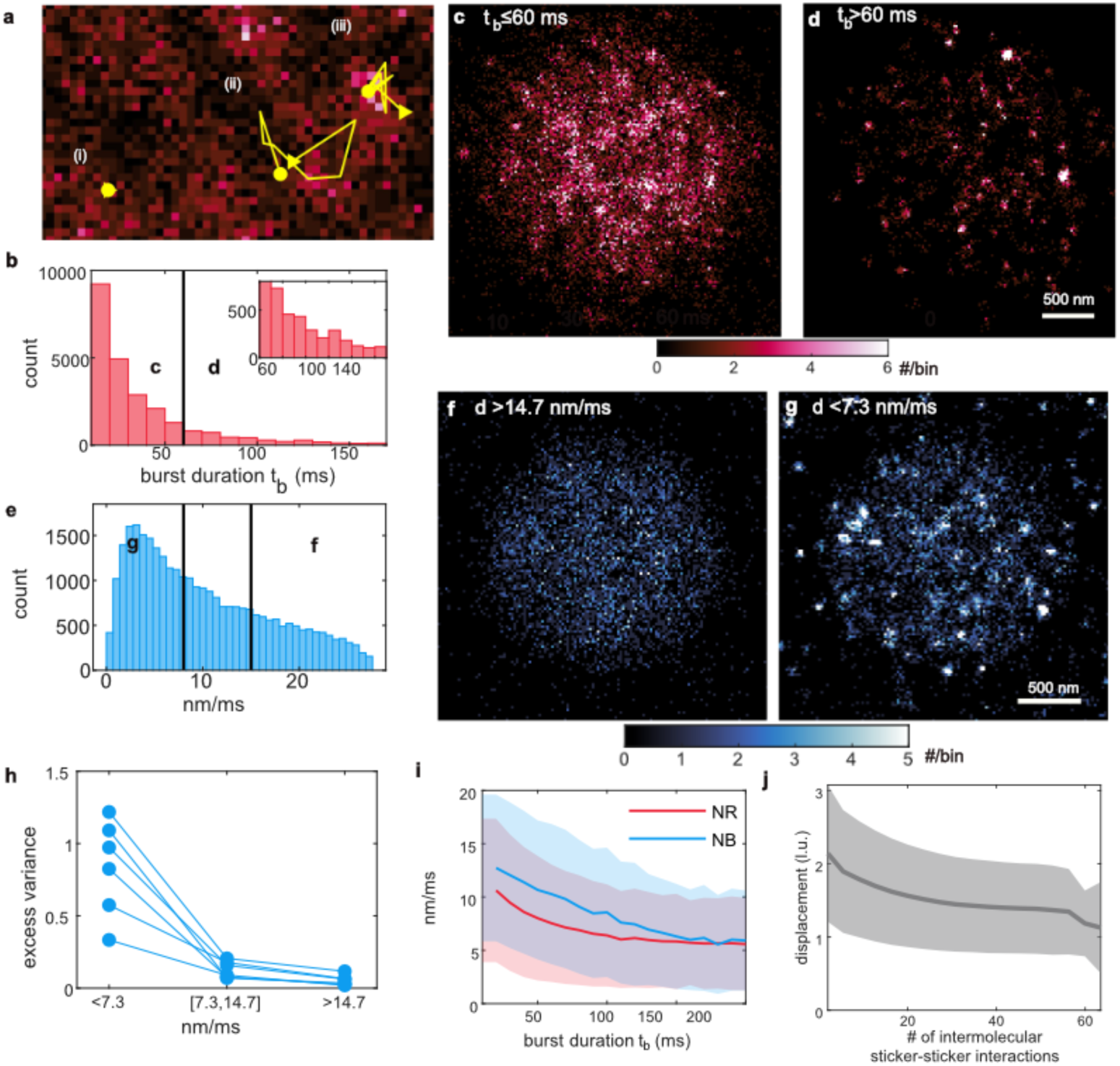
Tracking of single-molecule fluorescence burst durations and speeds uncover inhomogeneous molecular organization and dynamics within condensates. **a:** Single NR fluorophores exhibit both (**i**) short and (**ii, iii**) long burst durations, as well as (**ii**) high and (**iii**) low speeds. **b:** burst duration t_b_ of NR. **c, d:** SMLM images of NR molecules with burst durations (**c**) shorter than 60 ms and (**d**) longer than 60 ms. **Extended data** Fig. 5 **j-k** shows burst duration data for NB. **e:** Speed distribution of NB measured between consecutive camera frames (10 ms exposure time). **f, g:** SMLM images of NB molecules with speeds (**f**) larger than 14.7 nm/ms and (**g**) smaller than 7.3 nm/ms. **Extended data** Fig. 5 **e-g** shows speed data for NR. **h:** Excess variance of NB localizations grouped by speeds (left: <7.3 nm/ms, middle: between 7.3/ms and 14.7 nm/ms, right: >14.7 nm/ms). Blue lines: excess variance of NB within each condensate. **i:** Speeds of NB (blue) and NR (red) as a function of their fluorescence burst durations. Lines: mean value averaged over 150,000 trajectories for NB and 129,000 trajectories for NR; shaded region: ±1 standard deviation. **j:** Displacement in lattice units (l.u.) of each chain within a simulated A1-LCD condensate as a function of the number of intermolecular sticker-sticker interactions. Results are from lattice-based Monte Carlo simulations ^13^.

We separated NB emitters into three categories based on their speeds that were measured between consecutive frames (10 ms exposure times, **Fig. 3e**). An SMLM image, reconstructed from high-speed emitters, shows a uniformly distributed localization density across the condensate (**Fig. 3f**). These fluorogens were localized within sufficiently hydrophobic environments to emit fluorescence but bind relatively weakly, resulting in relatively high speeds. An SMLM image reconstructed solely from low-speed emitters exhibits clusters and nonuniform localization patterns (**Fig. 3g**). This phenomenon is consistent across six different A1-LCD condensates where the average excess variance of single-molecule images reconstructed using emitters with low, medium, and high speeds of emitters are 0.84 ± 0.33, 0.13 ± 0.06, and 0.05 ± 0.04, respectively (**Fig. 3h**). Condensates imaged using NR showed similar trends (**Extended Data** Fig. 5e-5g). Overall, NR and NB emitters with longer burst durations tend to have lower speeds (**Fig. 3i**). The hubs revealed by longer-burst emitters and those sensed by slowly moving emitters are consistent with one another (**Fig. 3** and **Extended Data** Fig. 5).

To provide a physical interpretation for why NR and NB dyes are trapped for longer periods of time at hubs, we analyzed published results from lattice-based LaSSI simulations ^13^. PLCDs are linear associative biopolymers featuring cohesive motifs known as stickers that are interspersed by solubility-determining spacers ^49^. Interactions for A1-LCD and related PLCDs follow a hierarchy whereby interactions between aromatic residues are the strongest ^13^. Stickers form reversible physical crosslinks with one another, whereas spacers modulate sticker-sticker interactions and the coupling between phase separation and percolation ^49, 50^. Aromatic residues of A1-LCD and related PLCDs function as stickers that enable physical crosslinking of these molecules ^2, 28^. Conversely, charged, and polar residues interspersed between the stickers act as spacers ^2, 28^. The simulations of Farag et al.^13^ showed that the spatial inhomogeneity and network connectivity within PLCD condensates are governed by the valence of stickers ^28^ and differences in interaction strengths between different stickers and spacers ^13^.

We propose that the nanoscale hubs uncovered using NR and NB are direct readouts of networked A1-LCD molecules formed via reversible, intermolecular physical crosslinks among aromatic stickers. Accordingly, we probed the correlation between molecular displacements and the extent of crosslinking in the simulated condensates. These simulations show that A1-LCD molecules that are part of densely crosslinked networks have smaller overall displacements (**Fig. 3j**). The simulations suggest that nanoscale hubs observed using single fluorogen imaging are clusters of A1-LCD molecules defined by the extent of physical crosslinking. The fluorogens become trapped for longer times in nanoscale hubs because of the higher local density of stickers within the clusters that underlie the hubs. Observations of dynamical inhomogeneities (**Fig. 3**) are a consequence of spatial inhomogeneities (**Fig. 2**). Similar reports for dynamical inhomogeneities arising from spatial inhomogeneities have been made by Shen et al.^51^ using adaptive single-molecule tracking in condensates that form at the post-synaptic density. Likewise, computations that reproduce structure factors from small-angle neutron scattering measurements also show spatial and dynamical inhomogeneities. Taken together with the work for Shen et al., ^51^ and Dar et al.,^52^ it appears that dynamical inhomogeneities within condensates are likely to be the signatures of spatial inhomogeneities that result from phase separation coupled to percolation ^50^.

### Valence of aromatic residues impacts nanoscale dynamics of PLCDs within condensates

The driving forces for phase separation of A1-LCD and designed variants thereof are governed by valence, i.e., the number and types of aromatic residues ^2, 28^. We used NR and NB dyes to obtain comparative assessments of nanoscale structures within condensates formed by two variants of the A1-LCD system designated as Aro^+^ and Aro^−^ ^28^. The Aro^+^ variant has more aromatic residues dispersed uniformly along the linear sequence when compared to the wildtype A1-LCD, while the Aro^−^ has fewer aromatic groups ^28^ (**Fig. 4a, Table S1**).

**Fig. 4.**
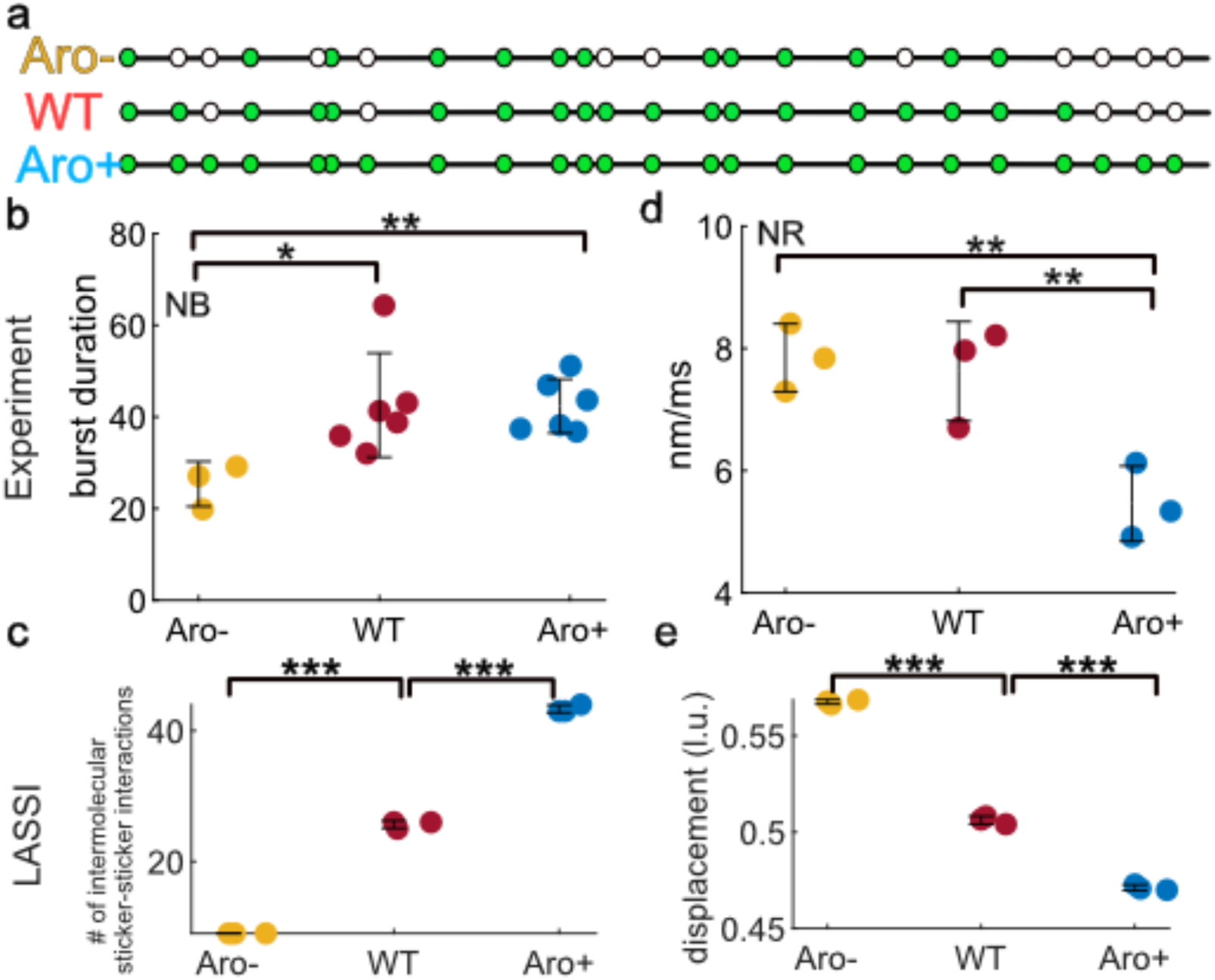
Nanoscale dynamics within condensates are influenced by the numbers of aromatic residues. **a:** Schematic showing the positions of aromatic amino acids as green circles in Aro^−^, A1-LCD (WT), and Aro^+^ variants. **b**: Fluorescence burst durations for the three condensates measured using NB. **c**: Number of intermolecular sticker-sticker interactions in simulated condensates. The sequence-specific simulations were performed using the LaSSI engine. **d**: Speed of NR within condensates formed by Aro^−^, WT, and Aro^+^. **e**: Displacements of protein chains quantified in simulated condensates. Circles in (b) represent the average burst durations of individual condensates, and in (d-e) they represent the median values of measurement parameters for individual condensates. Error bar: mean ± one standard deviation.

When probed using NB, the condensates formed by Aro^+^ demonstrated the longest burst duration with an average of 43 ms. In contrast, when probed using NB, the condensates formed by Aro^−^ show an average duration of 26 ms (**Fig. 4b**). Note that while dense phase concentrations of proteins change minimally within the condensates formed by different variants ^2, 13, 28^, the LaSSI simulations suggest that the extents of crosslinking vary with the valence of aromatic residues ^13^. The simulations show that the numbers of sticker-sticker interactions are highest in those formed by Aro^+^ and lowest in those formed by Aro^−^ (**Fig. 4c**). Based on these comparisons, we infer that the longer burst durations for NB in Aro^+^ condensates are due to the higher valence of aromatic residues. This larger valence results in higher extents of physical crosslinking and longer-lasting nanoscale hubs that trap NB for longer times and lead to longer fluorescence bursts. In contrast, condensates formed by Aro^−^ feature fewer sticker-sticker interactions, and this leads to lower affinities and shorter burst durations when the condensates are probed using NB. Similar data were obtained when the condensates were probed using NR (**Extended Data** Fig. 6).

These dynamics can be probed by measuring the speeds of fluorogen displacements. The measured speeds serve as useful proxies for evaluating how crosslinks, which generate locally elastic networks ^53^, impede the mobilities of fluorogens. Our single-molecule tracking measurements indicate that NR moves fastest in Aro^−^ condensates with an average speed of 8.9 nm/ms. In contrast, NR moves most slowly in Aro^+^ condensates with an average speed of 6.7 nm/ms (**Fig. 4d**). These observations indicate that the extent of crosslinking engenders restrictions to the movements of fluorogens. These observations are in line with findings from LaSSI simulations, which show that proteins in Aro^+^ condensates have the smallest displacements, whereas proteins in Aro^−^ condensates have the largest displacements (**Fig. 4e** and **Extended Data** Fig. 7).

### Single-molecule orientation localization microscopy reveals distinct features of interfaces

The fluorescence of MC540 is influenced by the equilibrium between dimer and monomer states of MC540, where monomers are fluorescent and dimers are non-fluorescent ^54^. This dimer-monomer equilibrium is strongly influenced by the molecular organization of the local environment ^55^. We leveraged polarized epifluorescence microscopy to probe the orientational preferences sensed by this fluorogen. We separate the fluorescence emission from MC540 into x- and y-polarized channels (𝐼*_x_*, 𝐼*_y_*). We then calculate the linear dichroism (LD) as the ratio: (𝐼*_x_* − 𝐼*_y_*)/(𝐼*_x_* + 𝐼*_y_*), which takes values from –1 to +1. An LD near +1 signifies that the x-polarization dominates the fluorescence emission, whereas a negative LD near –1 indicates dominance of the y-polarization. We noticed that transitioning from Aro^−^ to Aro^+^, the condensates exhibit increasingly discernible pink and green hues at the interfaces of condensates, indicating a stronger polarization preference at the interface (**Fig. 5a**). For Aro^+^, we observed mostly x-polarized fluorescence at the top and bottom edges and y-polarized fluorescence at the left and right sides (**Fig. 5a**). Comparing Aro^+^ with Aro^−^ and WT, we note that the LD distribution for Aro^+^ is broader and encompasses more instances with large absolute LD values (**Fig. 5b**).

**Fig. 5.**
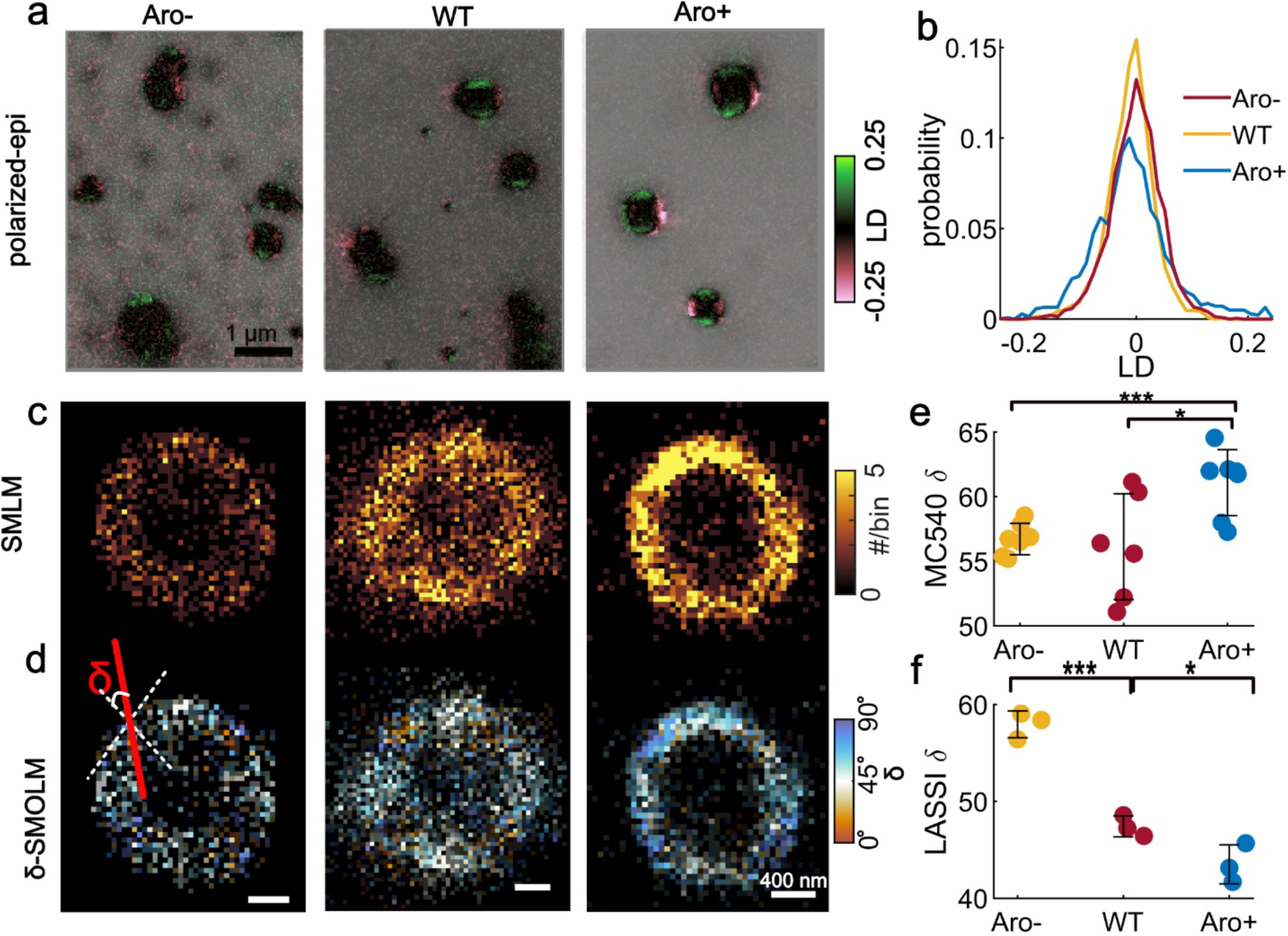
MC540 preferentially localizes to interfaces and displays distinct orientational preferences. **a:** Linear dichroism (LD) of MC540 measured by polarized epifluorescence imaging. **b:** LD distributions quantified from images shown in (**a**). **c:** SMLM images of MC540. **d:** Orientation angles δ for MC540 measured with respect to the normal vector to the condensate interface using SMOLM; the median angle δ is depicted within each 50 nm × 50 nm bin. **e:** Median δ values computed across individual condensates (circles). Error bar: mean±1 standard deviation. **f:** δ values computed from LaSSI simulations using orientations of protein molecules at the interface.

While SMLM images show a high density of MC540 at the interface of LCD condensates (**Fig. 5c**), they do not provide insights into how MC540 binds to the proteins. Thus, we employed single-molecule orientation localization microscopy (SMOLM) for measuring both the 3D position and orientation of MC540 (**Extended Data** Fig. 8 **and Movie S5**). SMOLM uses a pixOL phase mask to encode the information of 3D position and orientation into the images of single fluorogens ^56^. We quantified orientational preferences by quantifying the angle, denoted as δ, of MC540 in relation to the normal vector to the condensate interface (**Fig. 5d**). The SMOLM images, color-coded according to δ and referred to as δ-SMOLM, indicate that freely diffusing MC540 dyes have a statistically significant preference for orientations parallel to the condensate interface (depicted in bluish hues in **Fig. 5d**). This orientational preference is observed at the interfaces of all three PLCD condensates (**Fig. 5d**). When comparing the median δ values across multiple condensates for the three LCD variants, we observed that Aro^+^ exhibits the highest δ value, with a mean of 61°, while Aro^−^ and WT have similar mean values of δ, which are 57° and 56°, respectively (**Fig. 5e**).

In line with recent work, MC540 may be viewed as an adsorbent, whereas the PLCD molecules are the scaffolds ^57^. Erkamp et al., recently showed that scaffolds show statistical preference for orientations perpendicular to the plane of the interface, and adsorbents show a statistical preference for orientations parallel to condensate interfaces ^57^. Our SMOLM data for MC540 are in line with this expectation. Unlike the adsorbents, scaffolds should have a statistical preference for perpendicular orientations. This expectation was confirmed using results from LaSSI simulations, which show that scaffold molecules have statistically smaller δ values **(Fig. 5f** and **Extended Data** Fig. 9). The observed orientational preferences become more pronounced as the interactions among scaffolds become stronger with increasing numbers of aromatic residues **(Fig. 5f)**. Taken together with the simulation results, the SMOLM data, obtained using MC540, highlight the unique interfacial features of condensates.

## Discussion

The picture that emerges from single fluorogen imaging may be summarized as follows (**Fig. 6**): The interiors of condensates are more hydrophobic than the coexisting dilute phases. This is in line with recent inferences based on partitioning studies of small molecules that include drugs and drug-like molecules ^58^. Within condensates, we observe nanoscale spatial inhomogeneities that are manifest as nanoscale hubs, because the fluorogens, particularly NR, bind preferentially to hubs that are more hydrophobic than the background of the condensate. The nanoscale hubs have a greater ability to trap fluorogens within their hydrophobic pockets, thus enabling them to blink for longer times. Finally, SMOLM imaging suggests that the interface between dilute and dense phases is a unique environment where MC540 molecules show marked orientational preferences.

**Fig. 6.**
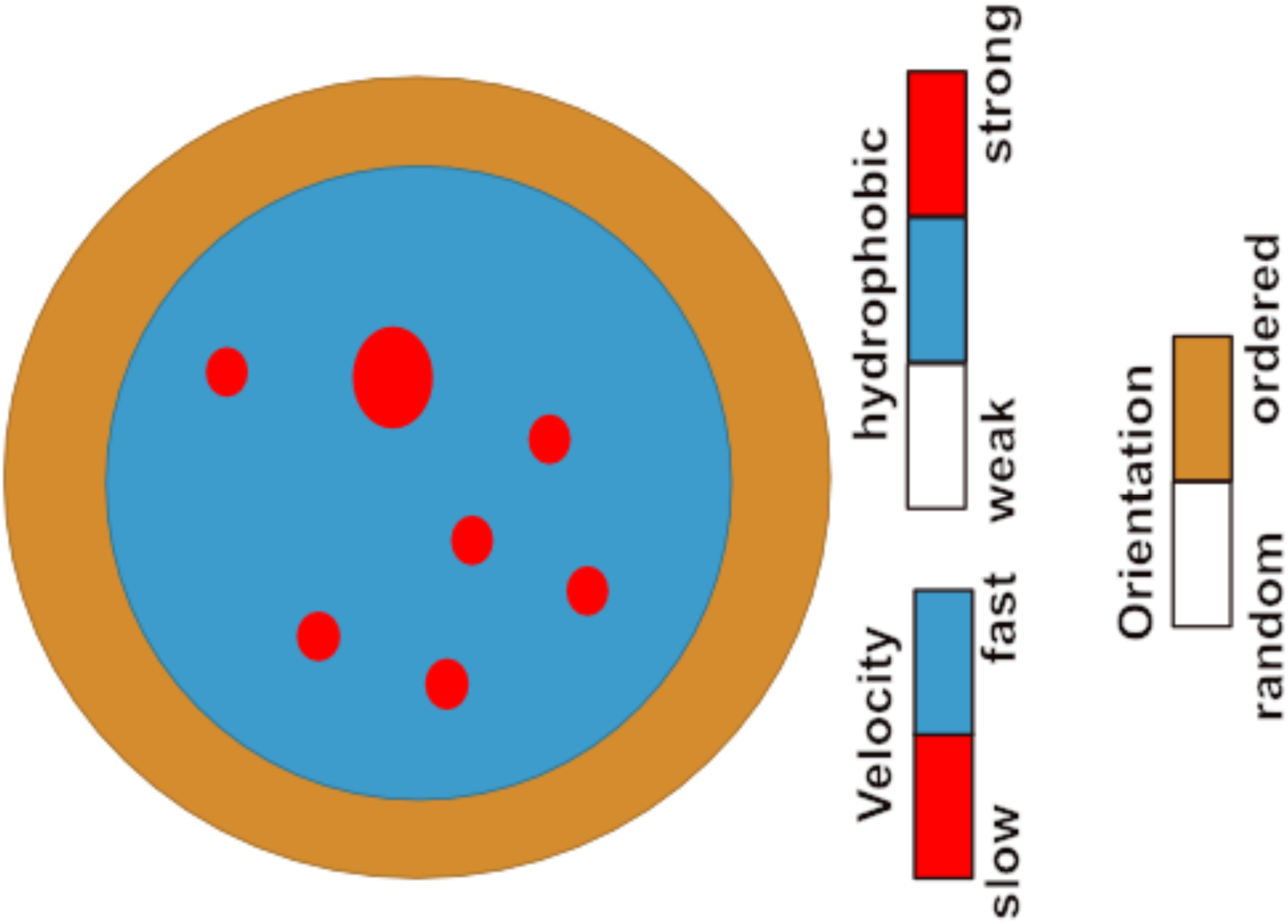
Schematic summarizing the structural features of condensates that were inferred from the fluorogenic experiments. Condensates are more hydrophobic than their coexisting dilute phases. They feature spatial inhomogeneities that are manifest as nanoscale hubs (red regions). Hubs are more hydrophobic compared to other regions (blue) within condensates, as well as to the dilute phase (white area). Fluorogens bound to nanoscale hubs move more slowly compared to other regions. Proteins at the interface (orange) have distinct orientational preferences that are unmasked by specific fluorogens such as MC540.

Other archetypes of low complexity domains that are drivers of condensate formation include RG-rich IDRs such as the N-terminal domain from the RNA helicase DDX4 ^3, 29^ (**Extended Data** Fig. 10). Although the molecular grammars of RG-rich IDRs are different from those of PLCDs ^13^, these domains appear to form condensates that respond in similar ways to NB and NR. This similarity is not surprising given recent findings regarding the π-character of arginine residues ^59^, and their apparent hydrophobicity ^60^ as probed by how water molecules organize around arginine sidechains ^61^. However, and in contrast to condensates formed by PLCDs and the RG-rich IDR of DDX4, the condensates formed by polynucleotides such as poly-rA respond very differently to NR (**Extended Data** Fig. 10). If there are spatial inhomogeneities and multiscale hubs within RNA condensates, they are not sensed by NR. The implication is that different types of fluorogens, specifically intercalating dyes, are needed to probe the internal organization of RNA-rich condensates ^57^.

We have presented the first structural characterization of inhomogeneities within condensates that leads to their descriptions as complex fluids with network-like structures ^52^. Inhomogeneities observed in simulations ^13^ been mapped onto graph-theoretic approaches for direct computations of condensate viscoelasticity ^10, 62^. We directly measured the predicted spatial and dynamical inhomogeneities within condensates. These observations help explain why condensates are viscoelastic materials ^5, 6, 7, 9, 10, 11, 30, 63, 64^. Continued development of new dye functionalities ^65, 66^, imaging hardware ^67^, and analytical tools ^68^ will help with uncovering spatiotemporal organization and dynamics within condensates, thus enabling direct assessments of structure-function relationships of biomolecular condensates.

The existence of inhomogeneities, albeit on the micron-scale, has been well established for multicomponent systems such as stress granules ^69^, nucleoli ^70^, nuclear speckles ^71^, the mitochondrial nucleoid ^4^, and synthetic systems comprising two or more macromolecular components ^72, 73^. Our work highlights the existence of nanoscale spatial inhomogeneities for condensates formed by a single type of macromolecule. We have shown how the varying stickers affects the observed inhomogeneities. The spatial inhomogeneities give condensates a sponge-like appearance. This is consistent with inferences made by Handwerger et al., who conjectured, based on quantitative analysis of differential interference microscopy images, that membraneless nuclear bodies including nucleoli, nuclear speckles, and Cajal bodies have sponge-like structures ^74^. Our findings suggest that multicomponent condensates that are characterized by micron-scale inhomogeneities might also feature nanoscale inhomogeneities. Generalizing our methods to characterize multicomponent systems is currently underway. To probe region-specific inhomogeneities, this effort will require functionalizing the fluorogens with peptide-based probes of molecular grammars that define distinct regions within larger, inhomogeneously organized multicomponent condensates.

## Acknowledgments

We thank J. Lu, N.A. Erkamp, and M-K. Shinn for help with the experiments. We thank T. Mittag and W.B. Borcherds for sharing their protocol for the expression and purification of the A1-LCD protein, and we thank L.E. Kay for sharing the DDX4 gene. This work was funded by the Air Force Office of Scientific Research grant (to RVP), the St. Jude Research Collaborative on the Biology and Biophysics of RNP granules (to RVP), and the National Institutes of Health (F32GM146418-01A1 to MRK and R35GM124858 to MDL). RVP and MDL are members of the Center for Biomolecular Condensates in the James McKelvey School of Engineering at Washington University in St. Louis.

## Author contributions

RVP and MDL conceived of and supervised the research. TW, MRK, and YQ designed and performed the imaging experiments and analyzed the data with inputs from MDL and RVP. MRK prepared A1-LCD, Aro^+^, Aro^−^, and DDX4 proteins, performed phase separation assays, and contributed intellectual insights. MF performed the LaSSI simulations and analyzed the simulation results. TW, MRK, MDL, and RVP wrote and edited the manuscript with input from all authors.

## Competing interests

RVP is a member of the scientific advisory board of and shareholder at Dewpoint Therapeutics Inc. The work reported here was not influenced by this affiliation. The remaining authors have no competing interests to declare.

## Additional information

Details of the imaging methods, analysis of images, constructs used, the preparation of condensates, LaSSI simulations, additional data and supplemental figures are presented in the supplementary material. Correspondence and requests for materials should be addressed to RVP and MDL.

---

**Extended Data Fig. 1.**
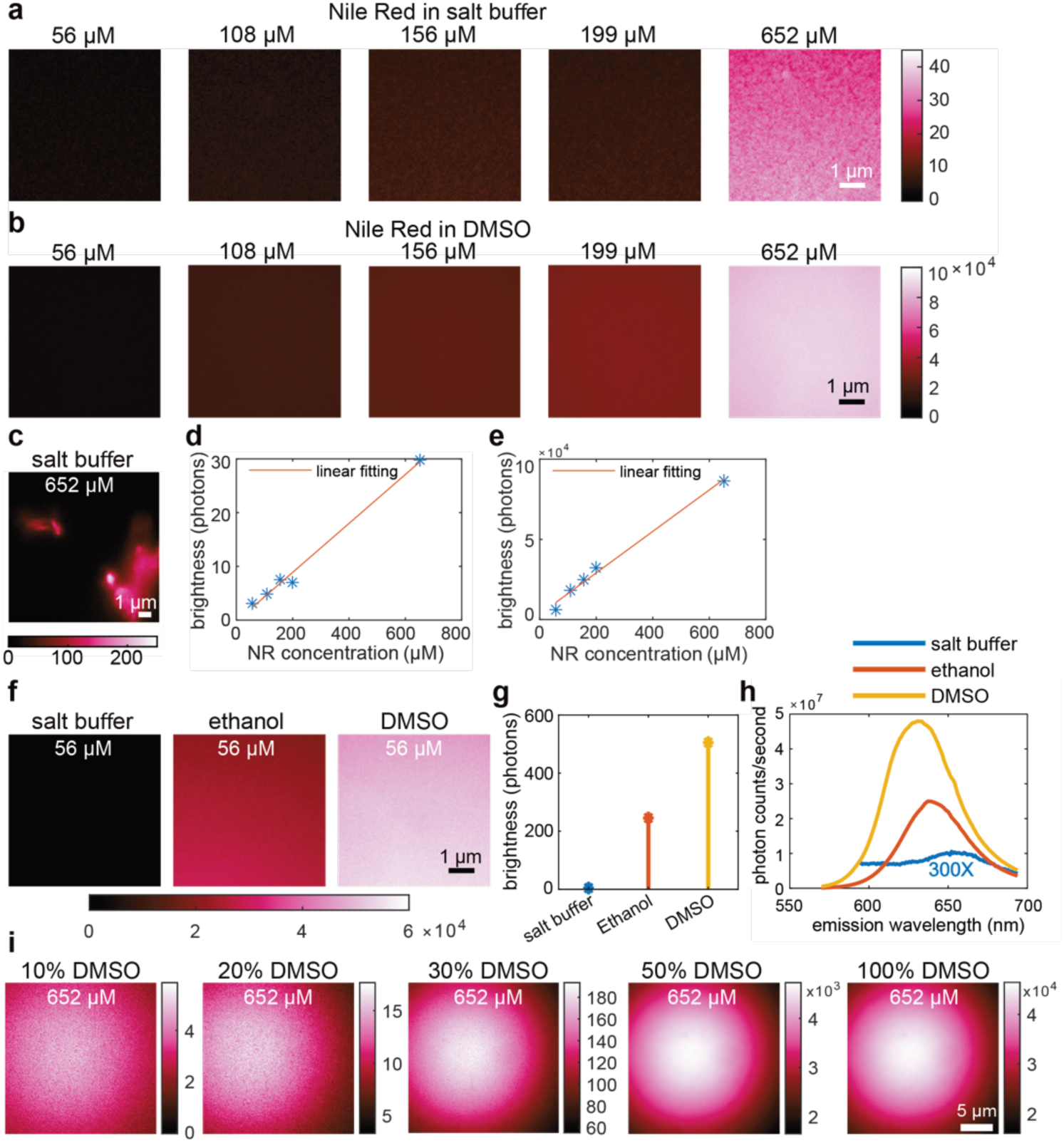
Calibration of the NR dye in homogeneous solutions. **a**: The intensity of NR at different concentrations dissolved in an aqueous buffer comprising 20 mM HEPES with 300 mM NaCl. This buffer is identical to the one used for preparing A1-LCD condensates. Images were captured with the microscope focused on the coverslip. Note that the intensity increases with NR concentration. **b**: Intensity of NR at different concentrations dissolved in DMSO. Images were captured on the coverslip. The intensity increases with NR concentration. Comparing panels in **a** to panels in **b**, we see that at equivalent concentrations of NR, the intensities in DMSO are at least three orders of magnitude higher than in aqueous buffers. **c:** In aqueous buffers, molecules of NR form aggregates at or above concentrations of 652 µM. This image is captured above the coverslip. The morphologies of NR aggregates are different from the hubs we observed in condensates. **d**: and **e:** There is a linear correspondence between brightness, quantified in terms of the number of photons collected, and the concentration of fluorogens we use, both in aqueous buffers (d) and in DMSO (e). **f:** Fluorescence intensity of 56 µm NR solutions in different solvents. These measurements highlight the increased brightness of the solvatochromic NR as the hydrophobicity of the solvent increases. Calibration of the increase in fluorescence intensities is provided by the color bar. **g**: The data in (**f**) are summarized by plotting the average intensities of the images. **h**: The emission spectrum of NR in three different solvents. The emission spectrum of NR in the aqueous buffer is scaled by a factor of 300 for better visualization. The emission spectrum of NR is blue-shifted as the hydrophobicity of the solvent increases. **i:** The dielectric constant of 100% DMSO is 46.7 and that of aqueous solutions is ∼78 at 296 K. At high concentrations of NR (652 µM), we quantified the fluorescence intensity as a function of % DMSO. These measurements show that the aggregates present in aqueous buffers (0% DMSO) are absent even in the presence of 10% DMSO. As the % of DMSO increases, the NR intensity increases by 2-4 orders of magnitude, thus highlighting the fluorogenic nature of NR.

**Extended Data Fig. 2.**
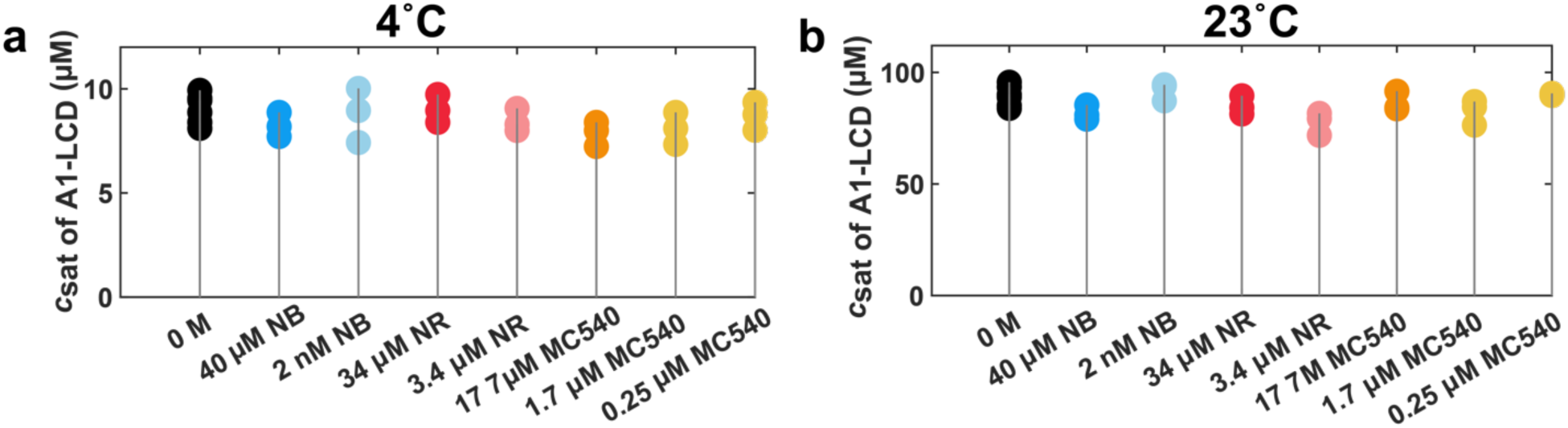
Quantifying the effects of different fluorogens on the driving forces for phase separation. Driving forces were quantified by measuring saturation concentrations (*c*_sat_) of A1-LCD molecules in the presence of different amounts of the different fluorogens used in this work. Within the error of the measurements, we find that adding NR, NB, and MC540 at different concentrations have minimal effects on the measured values of c_sat_. This inference is made based on comparisons to measurements made in the absence of dyes (black circles). All measurements were performed using a starting concentration of 150 µM A1-LCD in 20 mM HEPES buffer and 300 mM NaCl at (**a**) 4 °C and (**b**) 23 °C. Prepared protein and dye mixtures were allowed to phase separate for 30 minutes at the indicated temperatures before the dilute phase was separated from the dense phase via centrifugation and protein concentration measured using absorbance at 280 nm. Each set of samples was prepared in triplicate and measured independently; each triplicate measurement is shown as individual filled circles.

**Extended Data Fig. 3.**
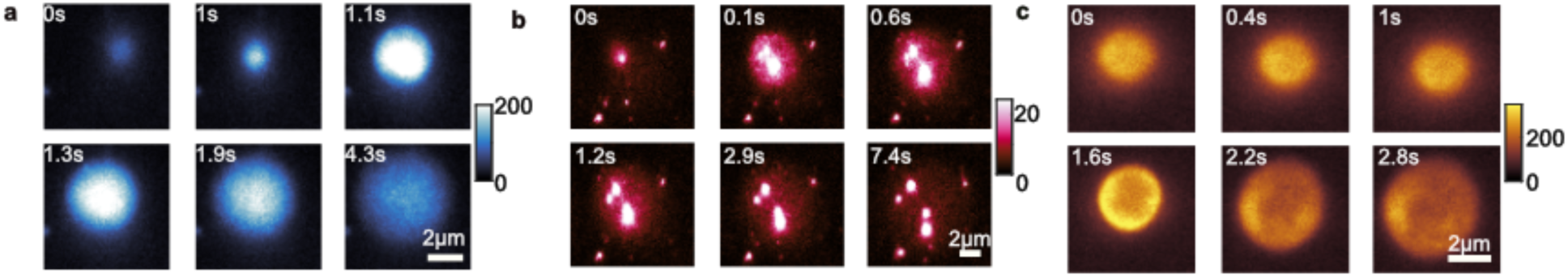
Dynamic wetting of the glass coverslip by A1-LCD condensates. The images were captured using (**a**) NB, (**b**) NR, and (**c**) MC540.

**Extended Data Fig. 4.**
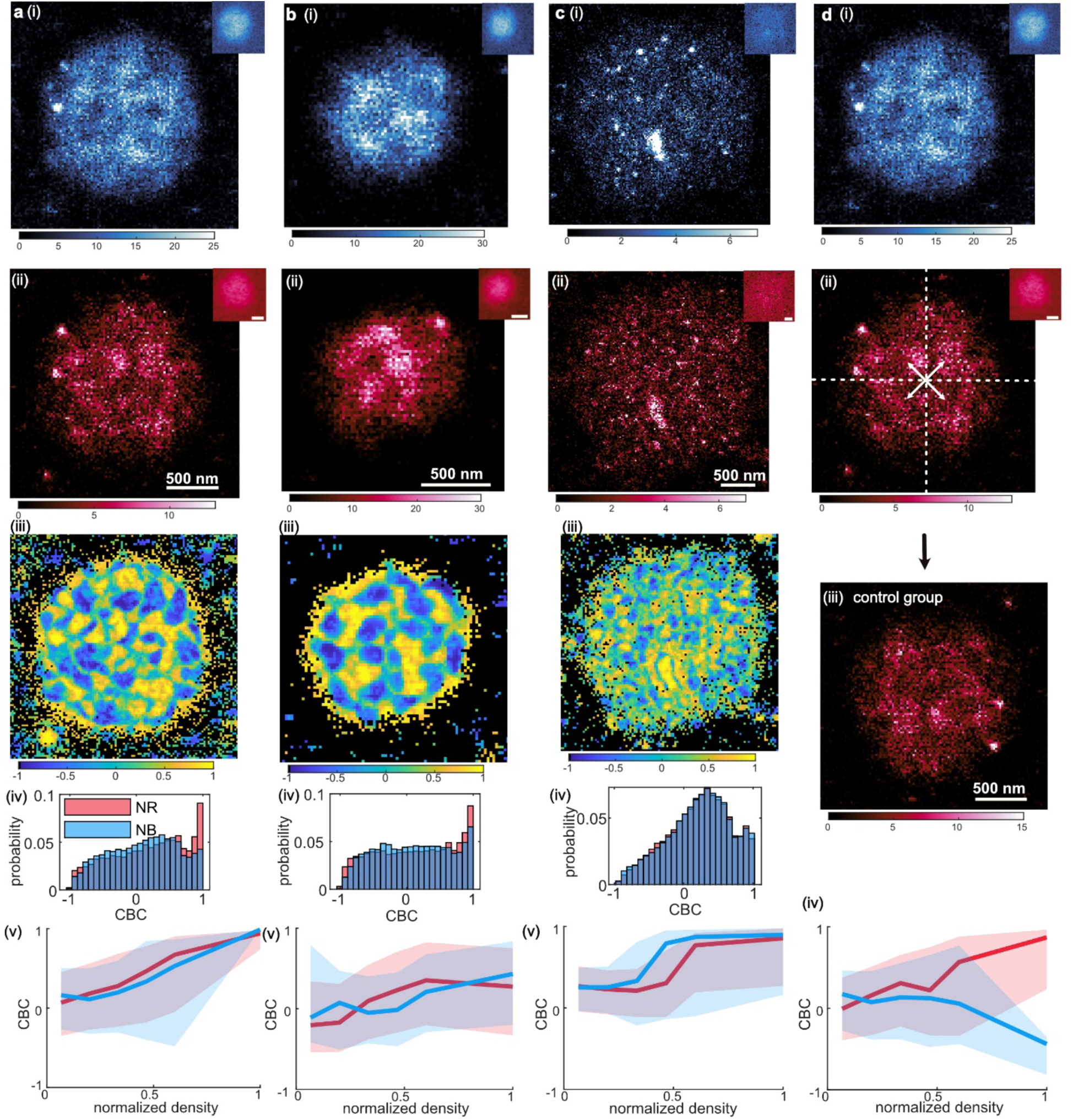
Spatial inhomogeneities within A1-LCD condensates imaged by NB and NR are highly correlated. **a**, **b**, and **c**: Correlation between NB and NR measured for three different A1-LCD condensates. SMLM images of single condensates collected using (i) NB and (ii) NR. Insets: epifluorescence images. Color bars: number of single molecules within each 20 nm × 20 nm bin. (iii) Coordinate-based correlation (CBC) between NR and NB localizations averaged for localizations within each 20 nm × 20 nm bin. (iv) Distribution of CBC scores for (blue) NB and (red) NR. (v) CBC of NB (blue) and NR (red) plotted as a function of localization density. Solid lines: median value; shaded area: 25^th^-75^th^ percentile. **d:** A condensate measured using (i) NB and (ii) NR as shown in **a**. We swapped emitters in the four regions of (ii) to form (iii) a modified NR condensate that should have weak correlations with the original condensate in **a**. (iv) CBC values between (i) NB localizations and (iii) the modified NR localizations.

**Extended Data Fig. 5.**
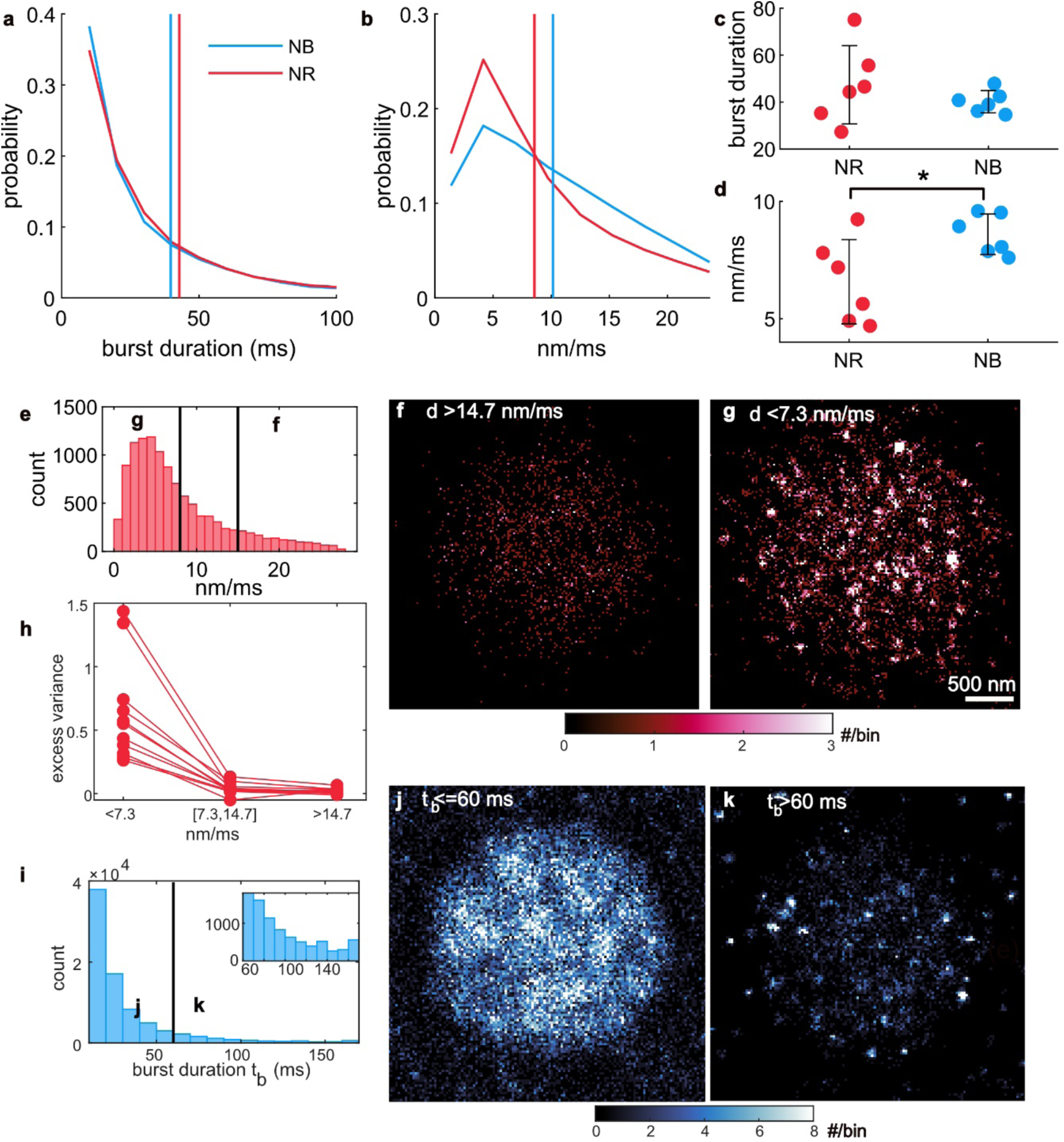
Tracking single-molecule fluorescence bursts and speeds reveal inhomogeneous molecular distributions within A1-LCD condensates. **a, b**: Distribution of (**a**) burst durations and (**b**) speeds for NB (blue) and NR (red). **c**: Mean burst duration for individual condensates. **d**: Median speed for fluorogens within individual condensates. Error bar: mean±1 standard deviation. Single-molecule tracking of (**e-h**) NR and (**i-k**) NB. **e:** NR speeds measured between consecutive camera frames (10 ms exposure time). **f, g:** SMLM images of NR with speeds (**f**) larger than 14.7 nm/ms and (**g**) shorter than 7.3 nm/ms. **h**: Excess variance of NR localizations grouped by speed (left: ≤ 7.3 nm/ms, middle: between 7.3/ms and 14.7 nm/ms, right: >14.7 nm/ms). Lines: excess variance of molecules within each condensate. **i:** Fluorescence burst durations t_b_ of NB. **j, k:** SMLM images of fluorogens (NB) with burst durations (**j**) shorter than 60 ms and (**k**) longer than 60 ms.

**Extended Data Fig. 6.**
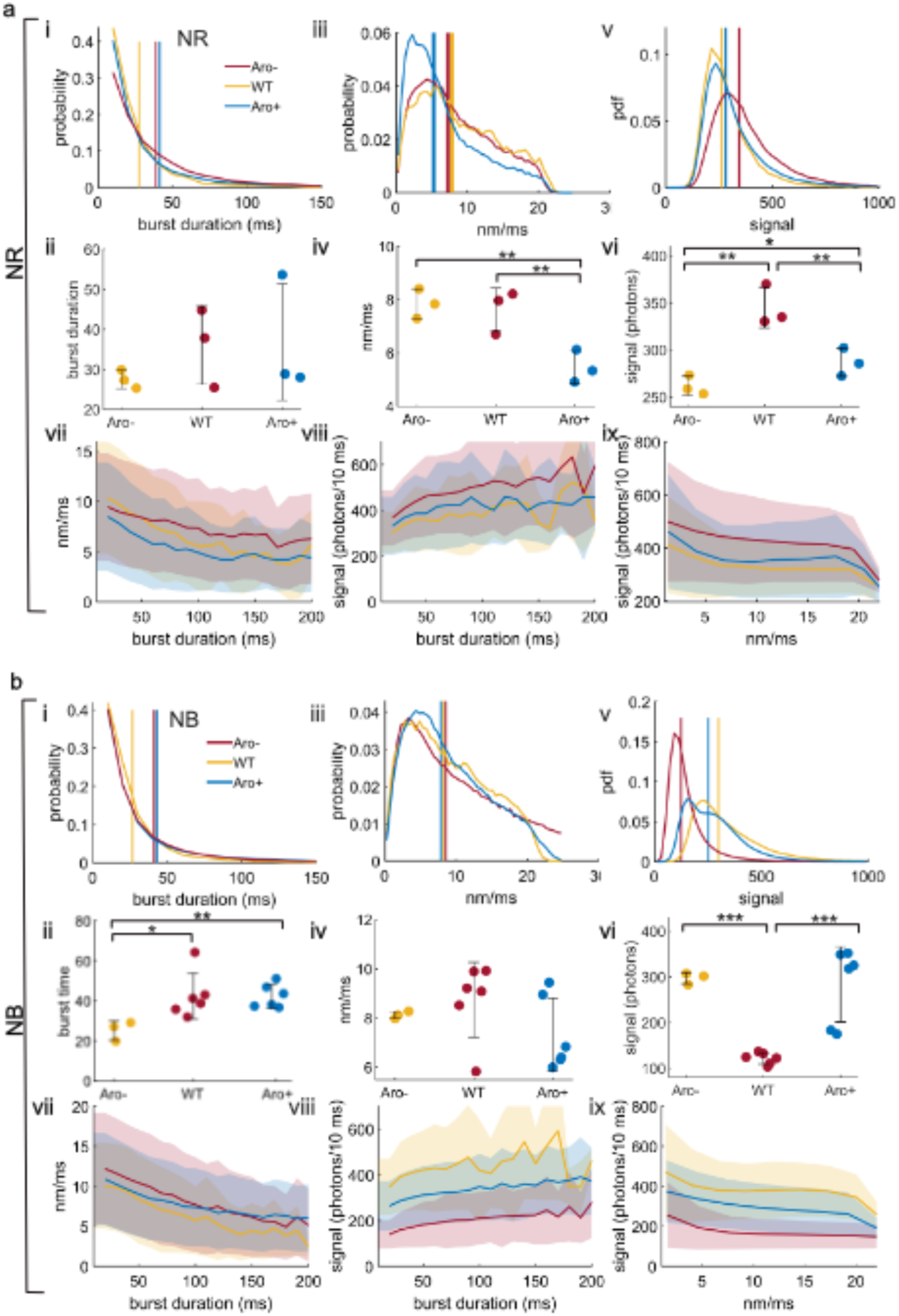
Speeds and burst durations of NR and NB molecules within Aro^−^, WT, and Aro^+^ condensates. **a**, **b**: Measurements of (a) NR and (b) NB for three different LCDs. **i,** Distribution of burst duration. Vertical lines: mean burst durations. **ii**, Mean burst duration for individual condensates. **iii**, Distribution of speed. Vertical lines: median speeds. **iv**, Median speed for individual condensates. **v**, Distribution of signal photons. Vertical lines: median signals. **vi**, Median signal of SMs in individual condensates. Error bar: mean±1 standard deviation. **vii**, Speeds of SMs as a function of their burst durations. **viii**, Signal photons of SMs as a function of their burst durations. **ix**, Signal photons as a function of their speed. Shaded region: ±1 standard deviation.

**Extended Data Fig. 7.**
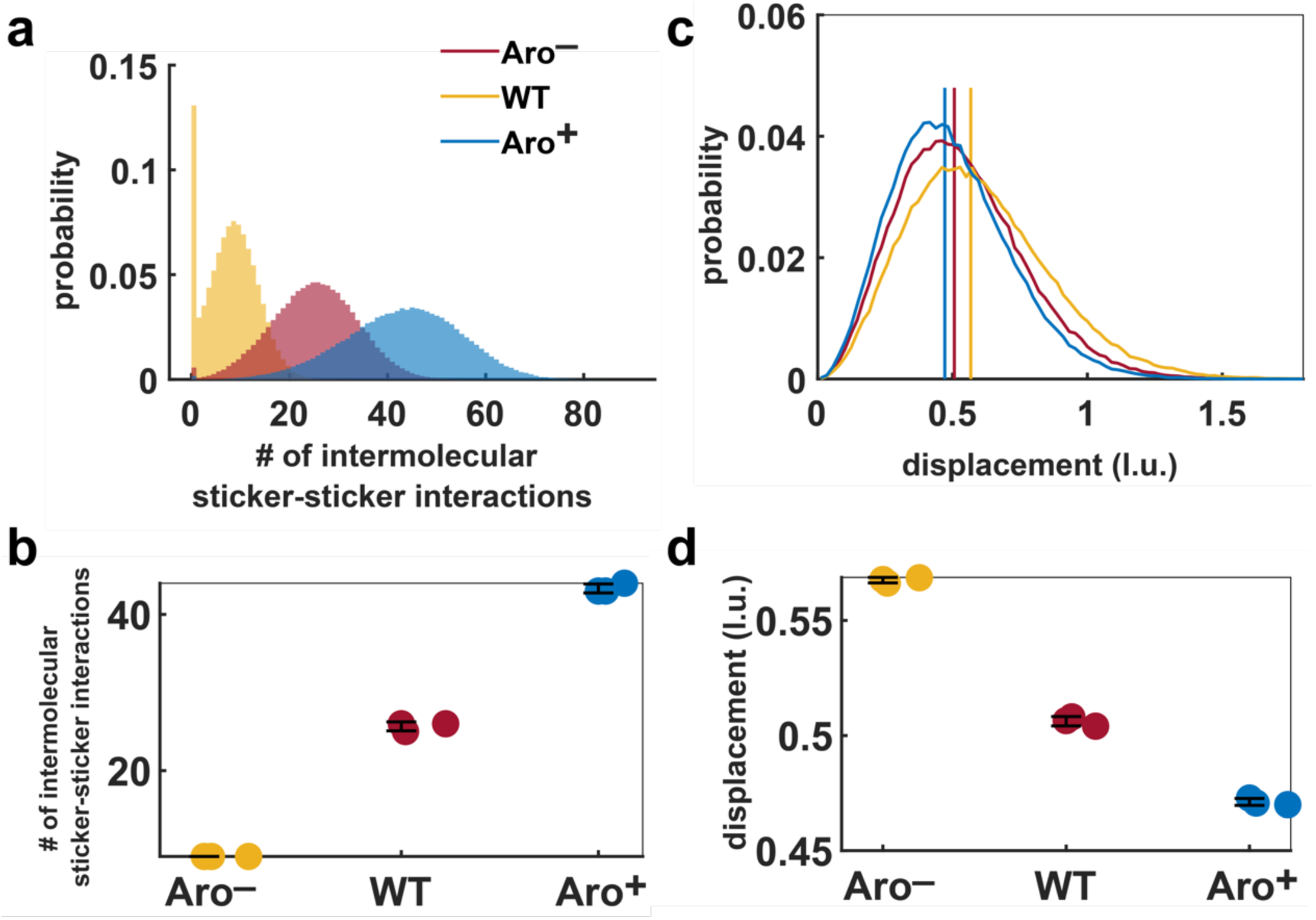
LaSSI simulations of condensates formed by Aro^−^, WT, and Aro^+^. **a**: Distribution of the number of sticker-sticker interactions. **b**: The median number of sticker-sticker interactions within individual condensates (reproduced from Fig. 4c). **c**: Distribution of displacement of protein chains. Vertical lines are the median displacements in lattice units (l.u). **d**: The median displacement of protein chains (reproduced from Fig. 4e). Error bars: mean±1 standard deviation.

**Extended Data Fig. 8.**
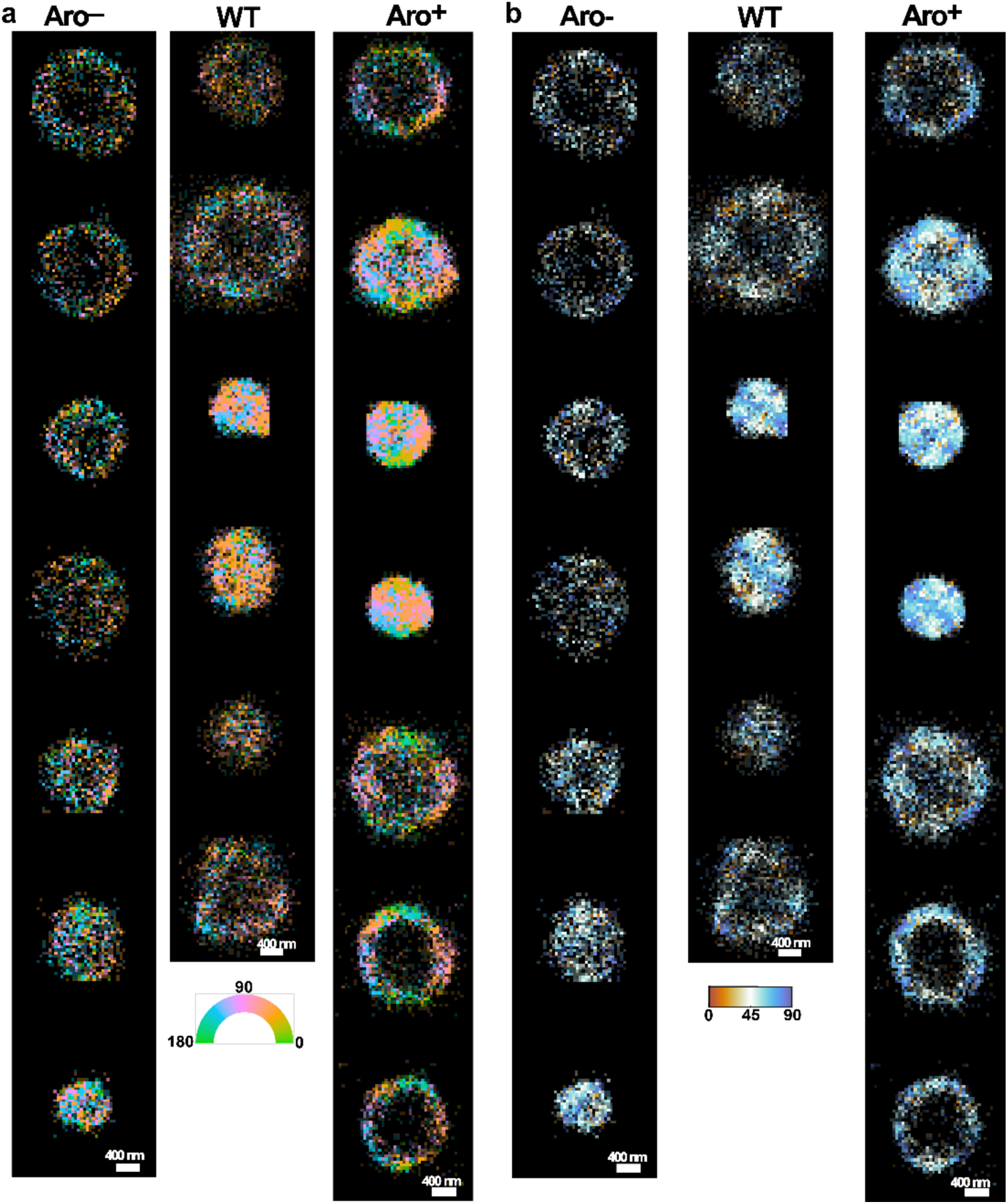
Single-molecule orientation-localization microscopy (SMOLM) of Aro-, WT, Aro+ condensates. **a**: Single-molecule orientation image with colors representing the measured azimuthal angles *φ* in the xy-plane. **b:** Single-molecule orientation image with colors representing the orientation angle *δ* measured with respect to the normal vector to the condensate interface. The images are reconstructed from localizations of freely diffusing MC540.

**Extended Data Fig. 9.**
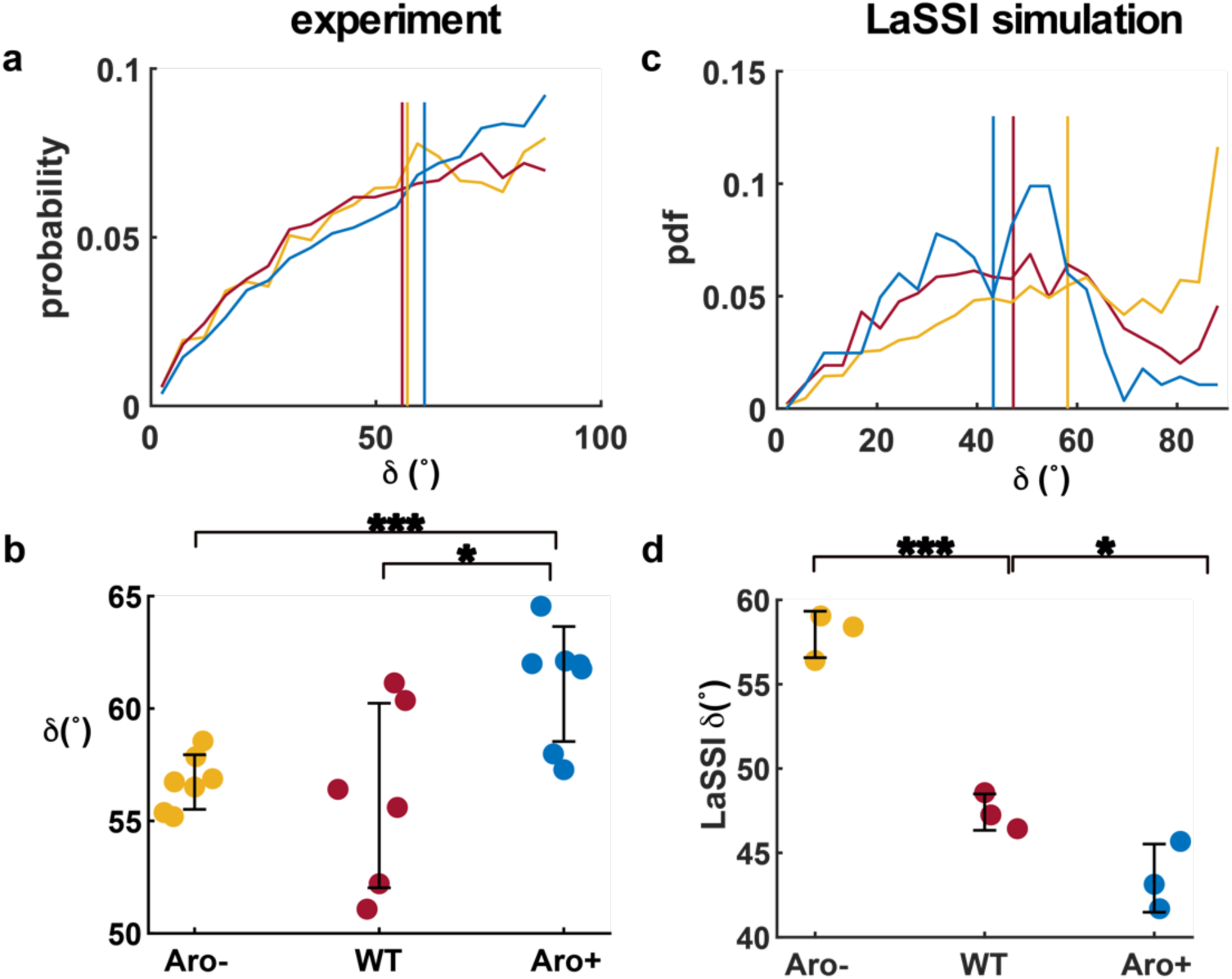
Orientation of MC540 measured by SMOLM and orientation of protein chains quantified in LaSSI simulations of different condensates. **a**: Distribution of the orientation angle *δ* for MC540 measured with respect to the normal vector to the condensate interface using SMOLM. Vertical lines: median value of orientation *δ*. **b**: The median orientation *δ* of MC540 within individual condensates (reproduced from Fig. 5e). **c**: Distribution of orientation *δ* of protein chains in LaSSI simulations. Vertical lines: median orientation *δ*. **d**: The median orientation *δ* of protein chains (reproduced from Fig. 5f). Error bar: mean±1 standard deviation.

**Extended Data Fig. 10.**
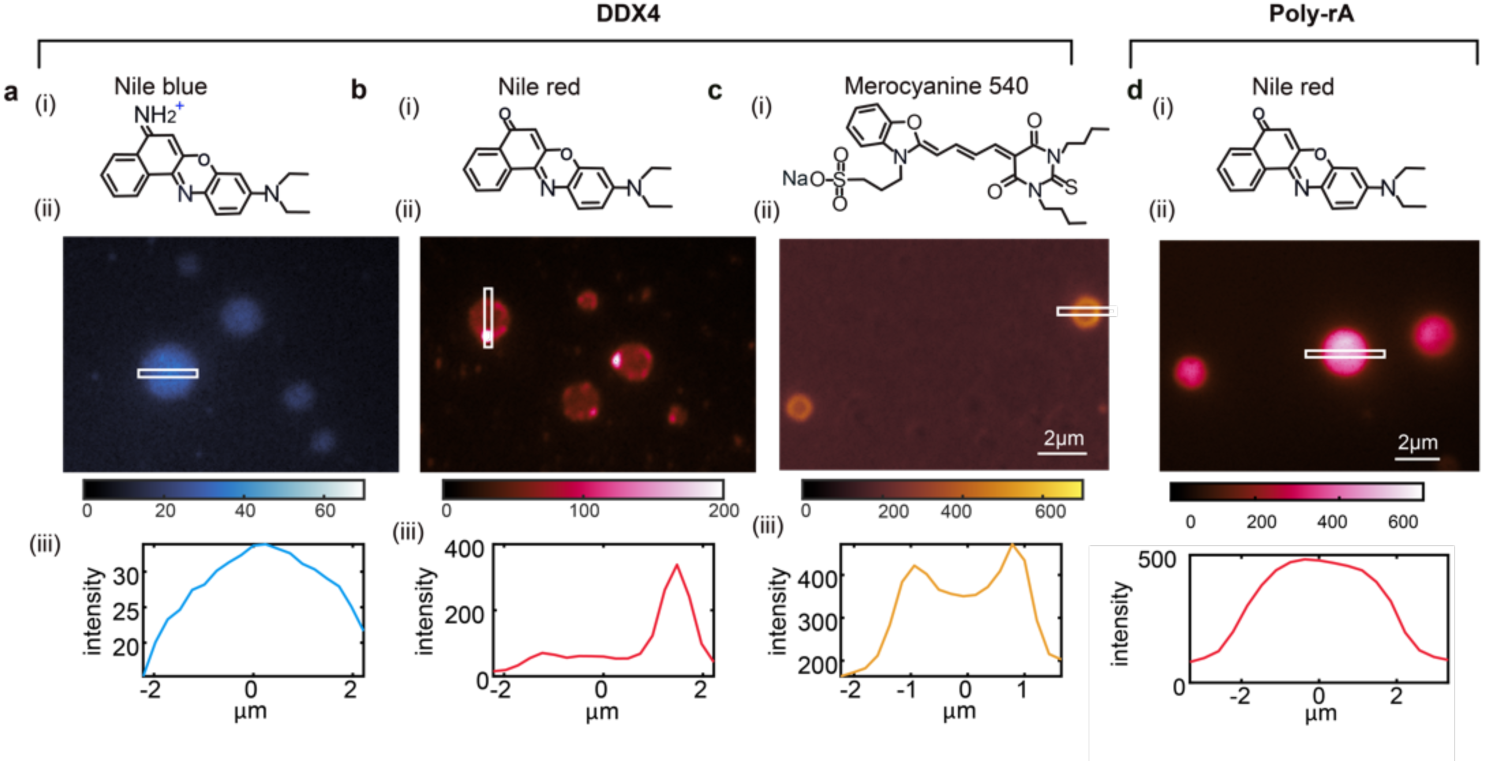
Epifluorescence microscopy reveals that different fluorogenic probes sense chemical environments inside DDX4-IDR condensates from different perspectives. Imaging DDX4-IDR condensates using **a.** Nile blue (NB), **b.** Nile red (NR), and **c.** merocyanine 540 (MC540). (i) Chemical structures. (ii) Typical condensates as viewed by epifluorescence microscopy. (iii) Fluorescence intensity profiles along the long axis of white box shown in (ii). **d**. Imaging poly-rA condensates using NR.

## Notes

https://osf.io/3qm2c/?view_only=5822386ed8704480a38509f79751d939

### Summary of Updates

Additional control/imaging experiments, more concise results and discussion in main text.

