## Supplementary Information for "Single fluorogen imaging reveals distinct environmental and structural features of biomolecular condensates"

### Materials and Methods

#### Sample Preparation

**Expression and purification of A1-LCD.** BL21-CODONPlus RIPL *Escherichia coli* cells (Agilent Technologies 230280) transfected with Δhexa_His-TEV-A1-LCD (either wild-type, Aro-, or Aro+) [1] were grown in LB broth (Sigma: L3522) in Erlenmeyer flasks with ≥ 5-fold head volume at 37˚C and 220 rotation per minute (RPM) orbital shaking until $OD_{600}$ ~0.6 was reached. Cultures were then chilled for 15 minutes in an ice bath before being induced with 0.35mM IPTG. Expression-induced cultures were grown for 6 hours at 37˚C. Cells were harvested by centrifugation, washed of residual media, flash frozen, and stored as pellets in 50 mL Falcon tubes at -80˚C.

At the start of purification, the cell pellet was gently resuspended to homogeneity in 35 mL supplemented lysis buffer (50 mM MES, 500 mM NaCl, 14.3 mM BME (β-Mercaptoethanol), 200 µM PMSF (phenylmethylsulfonyl fluoride) pH 6). Lysis buffer supplements are: 500U DNAaseI (Sigma - 4536282001), 500U RNAse A (Sigma - 10109169001), 5 mg Lysozyme (Sigma - 62971-10G-F), and one protease inhibitor tablet (Sigma - 40694200). The cell suspension was lysed via sonication on a Branson 550 sonicator with an L102C horn attachment using five series of the following 20-round cycle: 1 second on / 2 second off at 30% power. After a 30-minute spin at 38,000g, the pellet was resuspended in a resuspension buffer (6 M GdmCl, 20 mM Tris, 15 mM imidazole, 14.3 mM BME, and 200 µM PMSF; pH 7.5) via repeat pipetting and pulses of sonication until the solution was visibly homogenous. This solubilized pellet was spun at 38,000g for 30 minutes and the supernatant was applied to a gravity column with 5 mL bed volume of HiPur NiNTA resin (Fisher – 88223) equilibrated with resuspension buffer. The NiNTA resin was washed in 75mL wash Buffer (20 mM Tris, 30mM ImZ, 4 M Urea, 14.3mM BME, and 200 µM PMSF; pH 7.5), then protein was eluted in elution buffer (20 mM Tris, 350mM ImZ, 4 M Urea, 14.3mM BME, and 200 µM PMSF; pH 7.5). Peak fractions from NiNTA affinity purification were pooled and diluted 1:1 in dilution buffer (20 mM Tris and 14.3 mM BME; pH 7.5). To this, 1 mg of TEV protease was added, and the mixture was dialyzed overnight in dialysis buffer (20 mM Tris, 2 M urea, 50 mM NaCl, 0.5 mM EDTA, and 1 mM DTT; pH 7.5). The following morning, the protein solution was filtered through a 0.22 µm filter, then further purified via ion exchange chromatography on an ÄKTA Pure fast protein liquid chromatography (FPLC) module using a HiTrap SP 5mL column (Cytiva – 17115201). The column was equilibrated with buffer A (20 mM Tris, 2 M urea, 50 mM NaCl, and 14.3 mM BME; pH 7.5) and protein was bound then subjected to a continuous gradient protocol from 0.05M NaCl to 1 M NaCl. Fractions containing A1-LCD were pooled then further purified and buffer exchanged using size exclusion chromatography - HiLoad 16/600 Superdex 200pg column (Cytiva - 28989335) into the following storage buffer: 20mM MES and 4 M GdmCl at pH 5.5. Lastly, protein was pooled and concentrated in Amicon Ultra 3 MWCO (molecular weight cut-off) concentrator columns (Millipore-Sigma UFC 500396), according to the manufacturer's suggestions. Concentrated protein was stored at 4˚C until used.

**Expression and purification of DDX4-NT.** BL21 *E. coli* cells (NEB - C2530H) transfected with His-SUMO-DDX4-NT [2], a gift from Professor Lewis E. Kay, were grown in the same manner as A1-LCD. Cells were lysed in the same manner as A1-LCD, with the only difference being the supplemented lysis buffer (20 mM Sodium Phosphate, 750 mM NaCl, 20 mM Imidazole, 14.3 mM BME, and 0.2 mM PMSF; pH 7.5). The supernatant was recovered from a 25-minute spin at 38,000g and bound to an equilibrated HisTrap FF Crude 5 mL column (Cytiva

– 11000458) using an ÄKTA Pure fast protein liquid chromatography (FPLC) module. The NiNTA column was washed in 75 mL lysis buffer then protein was eluted in elution buffer (0.02 M Sodium Phosphate, 0.5 M NaCl, 0.35 M Imidazole, 0.0143 M BME, 0.0002 M PMSF; pH 7.5). Peak fractions from this affinity purification were pooled and diluted 5-fold in dilution buffer (0.02 M Sodium Phosphate, 0.0143 M BME; pH 7.5). This solution was further purified via ion exchange chromatography using a continuous gradient purification protocol with a HiTrap Heparin HP 5mL column (Cytiva – 17040703), Buffer A (0.02M Sodium Phosphate, 0.1M NaCl, 0.0143M BME; pH 7.5), and Buffer B (0.02 M Sodium Phosphate, 0.1 M NaCl, 0.0143 M BME; pH 7.5) on the ÄKTA Pure FPLC module. Peak fractions containing SUMO-DDX4 protein were pooled and cleaved of SUMO tags during an overnight dialysis in the presence of 0.02x ULP1 Protease in cleavage buffer (0.02 M Sodium Phosphate, 0.3 M NaCl, 0.001M DTT (Dithiothreitol), pH 7.5). DDX4-NT was purified to ≥98% using size exclusion chromatography - HiLoad 16/600 Superdex 200pg column (Cytiva - 28989335) on the ÄKTA Pure FPLC module in storage buffer (0.022M Sodium Phosphate, 1.1M NaCl, 0.0143M BME, pH 7.5). The DDX4-NT solution was supplemented with 10% glycerol and concentrated in Amicon Ultra 3 MWCO (molecular weight cut-off) concentrator columns (Millipore-Sigma UFC 500396) and concentrated protein was aliquoted into single-use volumes (typically 10 μL), flash froze in liquid $N_2$, and stored at -80˚C. Each step of the purification was assessed in the same manner as described for A1-LCD.

**Assessing protein purity.** After each stage of purification (namely affinity, ion exchange, size exclusion, and concentration steps), protein concentration and nucleotide levels were assessed on a Nanodrop2000 via absorbance at 280 nm measurements (A280) and A260/A280 ratiometric measurements. Similarly, after each purification stage, the inputs, flow-throughs, washes, and elutes were interrogated on using SDS-PAGE (sodium dodecyl sulfate–polyacrylamide gel electrophoresis) (Gel: BioRad miniPROTEAN TGX AnyKD - 4569036; MW latter: BioRad Precession Plus Protein Standard Unstained – 1610363), stained with EZblue Coomassie stain (10% (v/v) phosphoric acid, 10% (w/v) Ammonium sulfate, 20% (v/v) Methanol, 1.2% (w/v) Coomassie Blue) and de-stained via serial washes in $ddH_2O$. All proteins were purified to ≥ 99% purity assessed by densitometry of the Coomassie stained gels. Furthermore, all proteins exhibited an A260/A280 ratio of < 0.65, indicating no detectable nucleotide contamination.

**Preparation of condensates for imaging.** We adhered a silicone isolator (Grace Bio-labs, SKU 665206) to coverglass (Azer Scientific ES0107242, high precision 1.5H, 24×60 mm, 170±5 μm) to create a small chamber for imaging. To induce phase separation, we mixed 9 μL of ~100 μM A1-LCD (in 20 mM HEPES buffer, pH 7.0) with 1 μL of aqueous buffer (20 mM HEPES buffer with 3 M NaCl, pH 7.0) into the chamber to a final concentration of ~90 μM A1-LCD in 20 mM HEPES buffer with 300 mM NaCl. We then immediately added 0.5 μL of a solution (20 mM HEPES buffer with 300 mM NaCl, pH 7.0) containing a fluorogenic probe (final concentrations shown in **Table S2**) to the chamber. Phase separation occurs spontaneously within the chamber at room temperature (22° C) to form condensates for imaging. We noticed that condensates made with mixtures of His-tagged A1-LCD and untagged A1-LCD did not completely wet the coverslip, permitting condensates to remain distinct and unfused. We therefore use untagged A1-LCD for the wetting experiment shown in **Extended Data Fig. 3**, while an A1-LCD mixture containing ~20% His-tagged A1-LCD was used for experiments requiring distinct condensates shown in **Fig. 1, Fig. 2,** and **Fig. 3**. An identical procedure was followed for preparing Aro- and Aro+ condensates shown in **Figs. 4** and **5**, with the only difference being that we used a mixed 9 μL of ~160 μM Aro- A1-LCD to account for its lower saturation concentration ($c_{sat}$). To prepare DDX4 condensates for imaging, we mixed 1 µL of 2320 µM DDX4 (in 8 mM $NaH_2PO_4$, 12 mM $Na_2HPO_4$, 0.75 M NaCl) with 9 µL low-salt solution (8 mM $NaH_2PO_4$, 12 mM $Na_2HPO$, 0 M NaCl) into a small chamber for imaging.

**Preparation of condensates to measure driving forces for phase separation.** Driving forces were quantified by measuring saturation concentrations ($c_{sat}$) of A1-LCD molecules in the presence of different concentrations of fluorogens used in this work (**Table S2**). For each sample we induced phase separation, incubated the sample at a given temperature for 30 minutes, then separated the dilute phase from the dense phase via centrifugation at 14,000xg for 15 seconds. We then measured the protein concentration of the dilute phase. This is a standard assay for measuring saturation concentrations ($c_{sat}$) since the dilute phase concentration is the $c_{sat}$ that equalizes the chemical potential of the dense and dilute phases across the phase boundary. We mixed 42.5 µL of ~175 µM A1-LCD (in 20 mM HEPES buffer, pH 7.0) with 5 µL of salt buffer (20 mM HEPES buffer with 3 M NaCl, pH 7.0) and 2.5 µL of a fluorogenic solution (20 mM HEPES buffer with 300 mM NaCl, pH 7.0) all at once in a low protein retention Eppendorf tube (Fisher 11535564). The final concentration of A1-LCD was ~150 µM and the final concentration of dye was matched to those used in the imaging experiments (see **Table S2**); final solution conditions were 20 mM HEPES buffer with 300 mM NaCl. Each condition shown in **Extended Data Fig. 2** was prepared three times, in parallel. Prepared protein and dye mixtures were allowed to phase separate for 30 minutes at either 23˚C or 4˚C before the mixture was centrifuged, at its incubation temperature, at 14,000g for 30 seconds. 10 µL of the resultant dilute phase that was recovered, and protein concentration was measured using absorbance at 280 nm on a Nanodrop2000.

**Optical microscope setup and imaging procedure**

**Optical setup.** We used a home-built inverted microscope for epifluorescence imaging, single-molecule imaging, single-molecule tracking (**SI Fig. 1a**), polarized epifluorescence, and single-molecule orientation imaging (**SI Fig. 1b**). The microscope is equipped with an oil-immersion objective (OLYMPUS UPLSAPO100XOPSF, NA 1.4). The excitation laser, dichroic mirror, and emission filters were switched to match the excitation and emission spectra of NB, NR, and MC540 (**Table S3**). A relatively low peak laser intensity (~51 W/cm$^2$) and high concentration of fluorogens were used for epifluorescence imaging (**Table S2**). A relatively large peak laser intensity ~4088 W/cm$^2$ coupled with ultra-low concentrations of fluorogens facilitates bright fluorescence flashes for single-molecule imaging, tracking and orientation measurement.

**Single-molecule orientation localization measurement.** For measuring the orientation of MC540in condensates , we use the pixOL microscope which enables 3D position and 3D orientation measurement simultaneously[3] (**Fig. S1bc**). The pixOL microscope is designed to show different dipole-spread functions on the camera, depending on the orientation and 3D position of MC540 (**Fig. S1d-f**).

**Imaging protocols using multiple fluorogenic probes.** Since the emission and excitation spectra of MC540 and NB are well separated, we imaged single condensates using multiple probes by switching the excitation laser, dichroic mirror, and emission filter appropriately without crosstalk (**Table S3**). However, there is crosstalk between the emission spectra of NB and NR. The emission spectrum of NR significantly overlaps with the red window of our microscope (676±18 nm), while the emission of NB is barely observable in the orange window (593±23 nm, **Fig. S4**). Single NB molecules are virtually undetectable in the orange window (**Fig. S5**). To image a single condensate using both NB and NR, we always captured NB images first in the absence of

NR using the red-pass filter (**Table S3**), where the solution containing A1-LCD and NB were prepared according to the procedure above and **Table S2**. After NB imaging, we added 0.5 µL of a solution (20 mM HEPES buffer with 300 mM NaCl, pH 7.0) containing NR (final concentrations shown in **Table S2**) to the chamber. NR within the condensates was then imaged using the orange-pass filter. We observed stable consistent intensity values (epifluorescence imaging) and blinking dynamics (single-molecule imaging/tracking) for NR within seconds of adding it to the imaging chamber.

**Confocal imaging.** Point-scanning confocal microscopy was carried out on a LEICA STELLARIS 8 FELCON confocal microscope equipped with a white-light laser and adjustable band pass emission filters. This set up allowed imaging under the same excitation and emission wavelengths as was used in the custom-built microscope (see parameters for Nile Red and Nile Blue in **Table S3**). Samples were prepared as described in **Preparation of condensates for imaging** and concentrations of dyes matched to those used for epifluorescence imaging (see **Table 2**). These samples were imaged using a 100x Oil-immersion HC PL APO objective (1.40 NA) and a Leica Power HyD R detector. Z-stacks were taken in 0.5µm intervals from the coverslip surface up to at least 7 µm above this point**.** Images of stacks were taken sequentially using DIC/brightfield illumination followed by confocal imaging and at 8-bit depth. These image stacks were saved as TIFFs and analyzed using ImageJ (see **Data Analysis** section).

#### Data analysis

**Single-molecule detection and estimation algorithm.** SMLM depends on sequential detection and localization of individual molecular blinking events. These events are stochastic, and the improvement of temporal resolution has been enabled by the work of Mazidi et al.,[4] who developed a novel algorithm for analyzing frames with a high density of active molecules, or molecules whose images overlap. This approach enables accurate measurements of locations. Mazidi et al., showed that the spatial distribution of localization errors within super-resolved images tend to be vectorial in nature, and this can lead to systematic biases that degrade the resolutions of images reconstructed from blinking events. Mazidi et al., also showed that the shape of the point-spread function of the microscope has a fundamental effect on imaging artifacts. The algorithm of Mazidi et al., which we use in our work, is known as Robust Statistical Estimation algorithm (RoSE). This algorithm minimizes biases by estimating the likelihood of blinking events to localize molecules more accurately and eliminate false localizations. We used the bespoke regularized maximum likelihood estimator RoSE [4] to estimate single-molecule positions, brightness, and fluorescence burst durations. A regularization parameter of 0.3 and a calibrated point-spread function [5] model was used for estimation to account for optical aberrations. Details of the RoSE algorithm have been published by Mazidi et al. and are not reproduced here.

**Single-molecule orientation estimation.** Single-molecule orientation measurement requires detection of fluorescence "flashes" within raw camera images and estimating their 3D positions and orientations based on the shape of "flashes". We use a bespoke maximum likelihood estimator, RoSEO3D, to estimate the orientation and 3D position as described by Wu et al., [3].

**Calculating the orientation angle $\boldsymbol{\delta}$.** To calculate the orientation angle for MC540 measured with respect to the normal vector to the condensate interface, we first fit the estimated 3D condensate structure to an ellipsoid (**Fig. S7**). The normal vector to the condensate interface is calculated based on the surface of fitted ellipsoid. Next, we calculate the angle $\boldsymbol{\delta}$ between the estimated orientation $[\boldsymbol{\theta}, \boldsymbol{\phi}]$ and the surface normal. For comparing the orientation $\boldsymbol{\delta}$ among three A1-LCD variants shown in **Fig. 5e**, we only consider emitters with high estimation precision, namely emitters with a signal count larger than 500 photons, with polar angle $\boldsymbol{\theta}$ larger than $\mathbf{60}°$, and with axial positions h in [100, 800] nm. These filters ensure precise and reliable measurements of $\boldsymbol{\delta}$, since dim molecules high above the coverslip are more difficult to measure.

**Single-molecule tracking analysis – grouping localizations and quantifying burst durations and displacement trajectories.** At low concentrations of fluorogens, emitters are sparse enough to be easily distinguishable from one another. Two localizations are classified as originating from a single emitter if they appear within two adjacent camera frames and are separated by a distance smaller than a threshold T = $175 + 4\sigma_r$ nm, where $\sigma_r$ is the localization precision that varies for emitters with different signal-to-background ratios (~ 15 nm on average). Localizations collected at 10 ms intervals (the exposure time of the camera) are grouped together into a single trajectory until an emitter becomes dark, namely, when all emitters in the next frame are separated by a distance > T from the current emitter. The burst duration of a fluorescence emitter is equal to $N_f \times 10$ ms, where $N_f$ is the number of frames in the trajectory (**Fig. 3b**). The speed of an emitter is calculated as the Euclidean distance between its positions in each pair of consecutive frames within a trajectory (**Fig. 3e, h, i**); if the trajectory is longer than two frames, then each speed, calculated from a pair of frames, is reported separately (e.g., in **Fig. 3e**).

**Quantifying the degree of clustering within each condensate.** The clustering coefficient is calculated for each localization based on Ripley's $H(r)$ and Getis and Franklin's $L(r)$ functions [5,6]. To assign a cluster coefficient to the $i^{th}$ localization $p_i$, we define a circular region centered at $p_i$ with a radius $r$ of 50 nm. The radius $r$ is chosen to be larger than the localization precision, ~15 nm, of SMLM. This choice also ensures against blurring out the local information. The number of single molecules within this circular region is used to quantify:

$$H(r) = \sqrt{\frac{N_{p_i}}{\lambda\pi}} - r \; ; (1)$$

Here, $N_{p_i}$ represents the number of SMs within the circular region and λ is the average localization density across each condensate. The cluster coefficient quantifies how clustered or disperse the measured distribution is when compared to a uniform distribution. Localizations are classified as clustered if $H(r)$ is above some threshold (e.g., 20 in **Fig. 2e**).

**Excess variance.** We note that if single molecule localizations are described by a homogenous Poisson process, otherwise referred to as complete spatial randomness, then we expect the mean localization density to match the variance of the localization density when both statistics are calculated across an entire condensate or multiple condensates. Thus, the heterogeneity of localizations within a condensate may be quantified using excess variance, defined as:

$$V = \frac{\text{variance}(S_{con})}{\text{mean}(S_{con})} - 1; (2)$$

Here, $S_{con}$ represents a collection of localization densities (per 20 nm × 20 nm bin) measured within a condensate. We expect a uniformly random distribution to have an excess variance of zero (**Fig. S3**). An excess variance larger than zero represents heterogeneous structures.

**Coordinate-based correlation (CBC) between fluorogens.** We quantified the colocalization between pairs of fluorogenic probes ($A$ and $B$) directly using the position coordinates provided by SMLM rather than the brightness of pixels within an image [8]. To calculate the CBC value for a specific localization $A_i$ from dataset $A$, the distribution of localizations from both species $A$ and $B$ around $A_i$ is calculated as:

$$D_{A_i,A}(r) = \frac{N_{A_i,A}(r)}{N_{A_i,A}(R_{max})} \frac{R_{max}^2}{r^2}; \ (3)$$

$$D_{A_i,B}(r) = \frac{N_{A_i,B}(r)}{N_{A_i,B}(R_{max})} \frac{R_{max}^2}{r^2}; \ (4)$$

where $N_{A_i,A}(r)$ and $N_{A_i,B}(r)$ are the number of localizations of species A and species B within the distance r around $A_i$ respectively. In our calculation, we set the $R_{max}$ to be 300 nm, which determines the size of the region over which correlations are calculated. We then calculate the rank correlation coefficient as:

$$S_{A_i} = \frac{\sum_{r_j=0}^{R_{max}} \left(O_{D_{A_i,A}}(r_j) - \bar{O}_{D_{A_i,A}}\right)\left(O_{D_{A_i,B}}(r_j) - \bar{O}_{D_{A_i,B}}\right)}{\sqrt{\left(\sum_{r_j=0}^{R_{max}} \left(O_{D_{A_i,A}}(r_j) - \bar{O}_{D_{A_i,A}}\right)^2\right)}\sqrt{\left(\sum_{r_j=0}^{R_{max}} \left(O_{D_{A_i,B}}(r_j) - \bar{O}_{D_{A_i,B}}\right)^2\right)}}; \ (5)$$

where $O_{D_{A_I,A}}$ is the rank of $D_{A_I,A}$, and $\bar{O}_{D_{A_I,A}}$ is the mean of $O_{D_{A_I,A}}$. The CBC value of $A_i$ is calculated using:

$$CBC_{A_i} = S_{A_i} \exp\left(-\frac{E_{A_i,B}}{R_{max}}\right); \ (6)$$

with the $E_{A_i,B}$ as the distance from $A_i$ to the nearest neighbor from species $B$. CBC values range from –1 to 1.

**Analysis of confocal images.** Condensates were imaged as a confocal Z-stack. All images shown are of a single Z-slice that is representative and 1.5 µm above the coverslip; brightness and contrast have been optimized. Line scans were carried out using the line tool and plot profile analysis in ImageJ. Line thickness was set to 3 pixels and the same length line was drawn through five condensates per condition (NR and NB). Condensates were analyzed if they had equivalent diameters. The resultant intensity profiles were used to obtain median and 95$^{th}$ percentile confidence interval values of dye intensity. Partition coefficients were determined as follows: using only Z-slices 1.5 µm above the coverslips, coordinates of condensates were calculated via Otsu thresholding. Condensates smaller than 30 pixels (0.1411 µm/pixel) were omitted. Using the analyze particle tool, the mean intensity value (in arbitrary units – AU) for individual condensates was saved as 'condensate signal'. To use these values to obtain partition coefficients (PCs), we carried out additional analysis as follows: An independent thresholding procedure was applied to duplicated images to obtain the aggregate intensity of all coordinates in the frame that fell outside the condensate signal which we term 'background'. Individual PCs are the difference of the condensate signal and background. PC values are reported as a normalized value where 1 is the background.

#### LaSSI simulations

Simulations were performed using LaSSI [9], a lattice-based Monte Carlo engine, as described previously [10]. Monte Carlo moves are accepted or rejected based on the Metropolis-Hastings criterion so that the probability of accepting a move is equal to min(1, exp($-\beta\Delta E$)), where $\beta = 1 / kT$. Here, $kT$ is the simulation temperature and $\Delta E$ is the change in total system energy associated with the attempted move. Total system energies were calculated using a nearest-neighbor model and previously derived interaction parameters, which have been shown to accurately capture the phase behavior of A1-LCD and variants thereof [1]. For each simulation, 200 distinct chain molecules comprised of 137 beads each were placed in a cubic lattice with a length of 120 lattice units. Previous calibrations have shown that the numbers of molecules used in these simulations are adequate to avoid problems due to finite size effects. The simulations were performed for $10^{12}$ Monte Carlo moves and were allowed to equilibrate such that the chains formed a single condensate with a coexisting dilute phase before any analysis was performed. Given our interest in the dynamics of the condensate chains, we limited the Monte Carlo move set to local moves, eliminating non-physical moves that are part of the total set of moves. The jumping distance of chains in the condensate was calculated by tracking the distance moved by the center of mass of each chain after $2.5\times10^8$ total system Monte Carlo moves. The number of sticker-sticker interactions was calculated by counting the number of intermolecular interactions between aromatic beads (Phe or Tyr). Beads are interacting if they are within $\sqrt{3}$ lattice units. We performed five independent simulations, and the results shown aggregate all replicates.

### Supplementary Tables

**Table S1. Protein sequences used in this study**. Amino acid sequences for the proteins used in this study: wild-type hnRNPA1 low-complexity-domain (A1-LCD WT), A1-LCD with reduced valance of aromatic residues (A1-LCD Aro -), A1-LCD with increased valance of aromatic residues (A1-LCD Aro +), A1-LCD with an N-terminal His-tag used for purification (A1-LCD with His-tag), and the RG-rich N-terminal IDR from DDX4 (DDX4-IDR). Color coding of amino acids is as follows: black – nonpolar (A, I, L, M, V), green – polar (G, H, N, S, T, Q, C), red – acidic (D, E), blue – basic (K, R), yellow – aromatic (F, W, Y), and magenta – proline (P). The amino acids left of the vertical line in A1-LCD with His-tag are part of the His-tag.

| | |
|---|---|
| A1-LCD WT | GSMASASSSQRGRSGSGNFGGGRGGGFGGNDNFGRGGNFSGRGGFGGSRGGGGYGGSGDGYNG<br>FGNDGSNFGGGGSYNDFGNYNNQSSNFGPMKGGNFGGRSSGGSGGGGQYFAKPRNQGGYGGSS<br>SSSSYGSGRRF |
| A1-LCD Aro - | GSMASASSSQRGRSGSGNSGGGRGGGFGGNDNFGRGGNSSGRGGFGGSRGGGGYGGSGDGYNG<br>FGNDGSNSGGGGSSNDFGNYNNQSSNFGPMKGGNFGGRSSGGSGGGGQYSAKPRNQGGYGGSS<br>SSSSSGSGRRF |
| A1-LCD Aro + | GSMAFASSFQRGRYGSGNFGGGRGGGFGGNDNFGRGGNFSGRGGFGGSRGGGGYGGSGDGYNG<br>FGNDGSNFGGGGSYNDFGNYNNQSSNFGPMKGGNFGGRSSGGSYGGGQYFAKPRNQGGYGGSS<br>FSSSYGSGRRF |
| A1-LCD with His-tag | MSYYHHHHHHLESTSLYKKAGFENLYFQ\|GSMASASSSQRGRSGSGNFGGGRGGGFGGNDNFG<br>RGGNFSGRGGFGGSRGGGGYGGSGDGYNGFGNDGSNFGGGGSYNDFGNYNNQSSNFGPMKGGN<br>FGGRSSGGSGGGGQYFAKPRNQGGYGGSSSSSSYGSGRRF |
| DDX4-IDR | MGDEDWEAEINPHMSSYVPIFEKDRYSGENGDNFNRTPASSSEMDDGPSRRDHFMKSGFASGR<br>NFGNRDAGECNKRDNTSTMGGFGVGKSFGNRGFSNSRFEDGDSSGFWRESSNDCEDNPTRNRG<br>FSKRGGYRDGNNSEASGPYRRGGRGSFRGCRGGFGLGSPNNDLDPDECMQRTGGLFGSRRPVL<br>SGTGNGDTSQSRSGSGSERGGYKGLNEEVITGSGKNSWKSEAEGGES |

**Table S2. The concentration of fluorogenic probes for epifluorescence imaging, single-molecule imaging, and tracking**.

| **Fluorogenic probes** | **Nile blue** | **Nile red** | **Merocyanine 540** |
|---|---|---|---|
| Epifluorescence imaging | 40 μM | 34 μM | 17 μM |
| Single-molecule imaging/tracking/orientation | 2 nM | 3.4 μM | 0.25 μM |

**Table S3. Microscope setup for imaging condensates using three fluorogenic probes**.

| **Fluorogenic probes** | **Nile blue** | **Nile red** | **Merocyanine 540** |
|---|---|---|---|
| Excitation laser | Coherent OBIS 637 nm | Coherent Sapphire 561 nm | Cobolt 06-DPL 532 nm |
| Dichroic mirror | Chroma ZT532/640rpc-UF1 | Semrock Di01-R488/561 | Chroma ZT532/640rpc-UF1 |
| Filter | Semrock FF01-676/37 | Semrock FF01-593/46 | Semrock FF01-593/46 |

**Table S4. Laser power used for pumping fluorogens**.

| **Imaging mode** | **Nile blue** | **Nile red** | **Merocyanine 540** |
|---|---|---|---|
| Epifluorescence/Confocal imaging | ~51 W/cm$^2$ | ~51 W/cm$^2$ | ~51 W/cm$^2$ |
| Single-molecule imaging/tracking/orientation | ~4088 W/cm$^2$ | ~4088 W/cm$^2$ | ~4088 W/cm$^2$ |

**Table S5. Fluorescence statistics of NB and NR**. The number of localizations for each condensate, fluorescence signal per localization, burst duration, and speed for measurements shown in **Fig. 3h** and **Extended Data Fig. 5h**. The NB data were averaged over six condensates and NR data were averaged over eleven condensates.

| | **Nile blue** | **Nile red** |
|---|---|---|
| Number of localizations/condensate | $6.0 \times 10^4$ | $2.9 \times 10^4$ |
| Average signal (photons/localization) | 149 | 173 |
| Average burst duration (ms) | 41 | 36 |
| Average speed of emitters (nm/ms) | 10 | 7.5 |

### Supplementary Figures

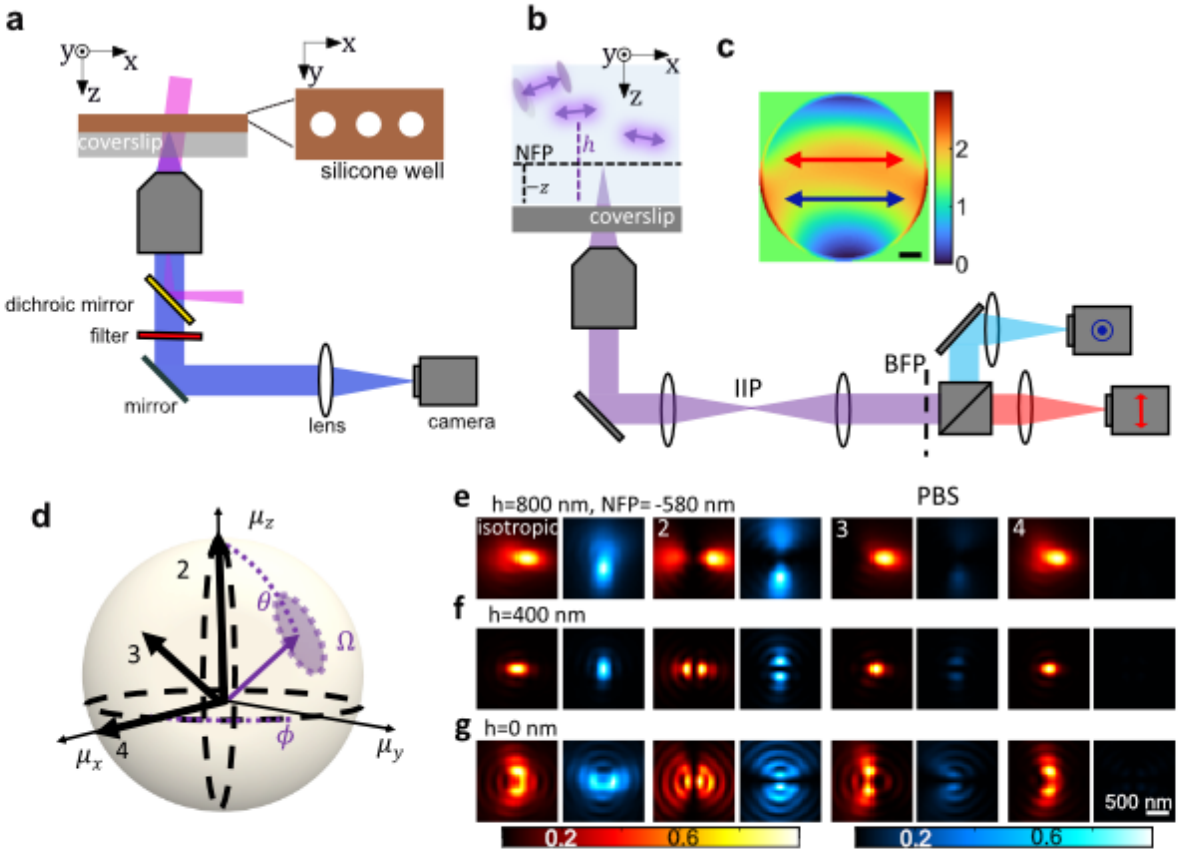

**Fig. S1. Schematic of the imaging system. a:** Solutions containing the condensates are deposited in silicone wells adhered to a coverslip. An excitation laser is coupled into an objective lens to illuminate the sample. Fluorescence from fluorogenic probes is collected from the focal plane of the objective and filtered by a dichroic mirror and a bandpass filter. After passing through a tube lens, the fluorescence is captured by a camera. **b**: For the polar-epifluorescence microscope used in **Fig 4**, a polarizing beam splitter is used to split the emission light into x/y-polarized channels and captured by two cameras or two areas of one camera. c: To measure the orientation of MC540 in **Fig 5**, we insert a pixOL phase mask shown in (**c**) into the back focal plane (BFP) of the polar-epifluorescence microscope shown in (**b**). **d**: Orientation of a dipole-like emitter, parameterized by a polar angle $\theta \in [0°, 90°]$, azimuthal angle $\phi \in (-180°, 180°]$, and a wobble solid angle $\Omega \in [0,2\pi]$ sr. **e-g**: Simulated images of emitters located at (e) h=800 nm, (f) h=400 nm and (g) h= 0 nm with orientations $[\theta, \phi, \Omega]$ shown in (d) [emitter 1: $\Omega = 2\pi$ sr, emitter 2: $(0°, 0°, 0)$, emitter 3: $(45°, 0°, 0)$, and emitter 4: $(90°, 0°, 0)$]. Emitters with different orientations and located at different positions will show dipole-spread functions with different intensity distributions.

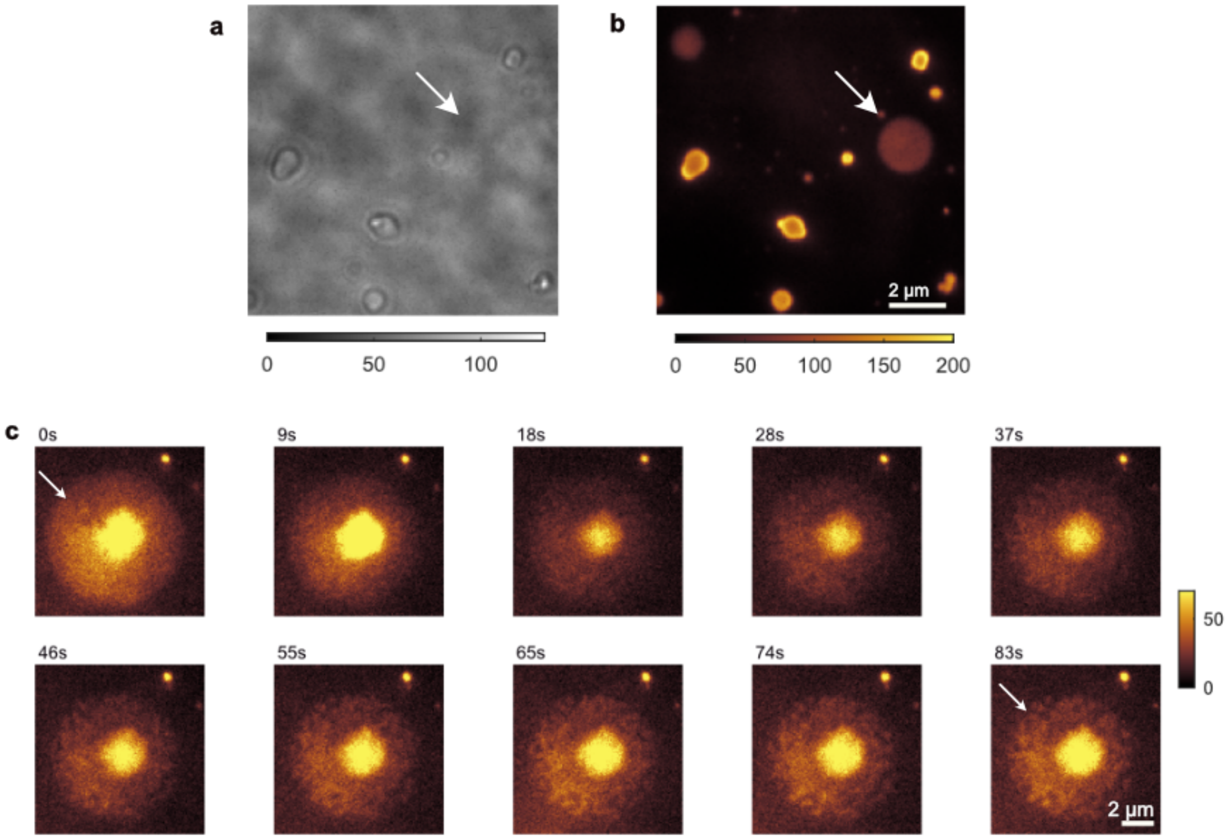

**Fig. S2. Imaging A1-LCD condensates using bright-field imaging and epifluorescence imaging of MC540. a:** Unwetted condensates have high contrast in the bright-field image. **b:** Images show high MC540 fluorescence at the interfaces of condensates that do not wet the coverslip. **c:** In contrast to the condensates that do not wet coverslips, the fully wetted condensates imaged at different times using MC540 show non-uniform structures (arrow) appearing within the condensates – compare the arrow at 0 s vs. 83s. Importantly, induced non-uniformities imaged by MC540 were only noticed in wetted condensates in the presence of high concentrations of MC540 (17 µM). Therefore, all analyses of interfacial features, specifically the analysis of orientational preferences quantified in **Fig. 5** used unwetted condensates and a low concentration of MC540 (0.25 µM) that avoids induction of artifacts while focusing on condensates that do not wet the coverslips.

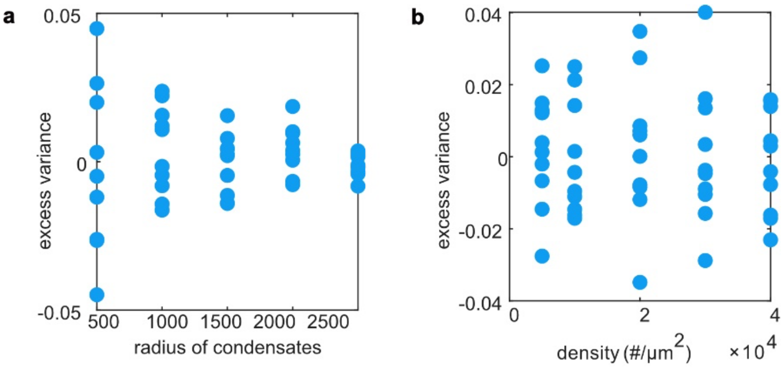

**Fig. S3. Excess variance is essentially zero for simulated homogeneous condensates with uniformly distributed localizations.** This trend holds true when the condensate radii are varied (**a**) and when the localization densities are varied (**b**). Ten simulations were performed for each condition.

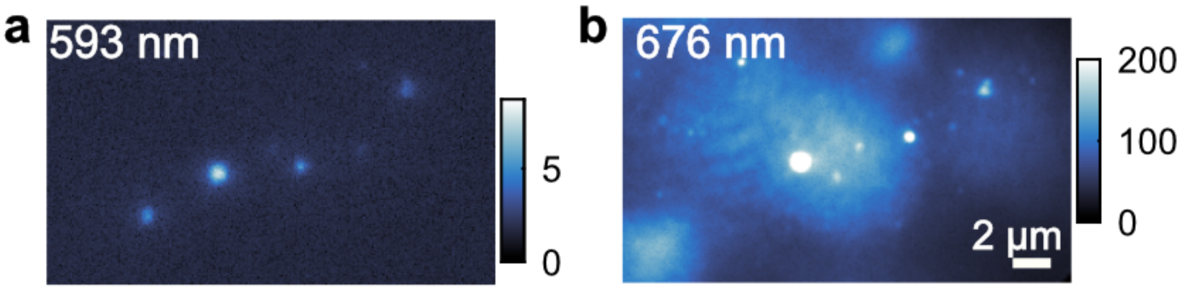

**Fig. S4. NB epifluorescence microscopy of A1-LCD condensates collected using two emission windows.** Fluorescence collected in the (a) orange emission window (593 nm) was excited using a 532-nm laser, and fluorescence collected in the (b) red emission window (676 nm) was excited using a 637-nm laser. The emission of NB is weak at the orange emission window based on the color bar in (a). Therefore, when imaging condensates using both NB and NR, capturing NB first will not significantly influence the signal in the NR channel (orange emission window).

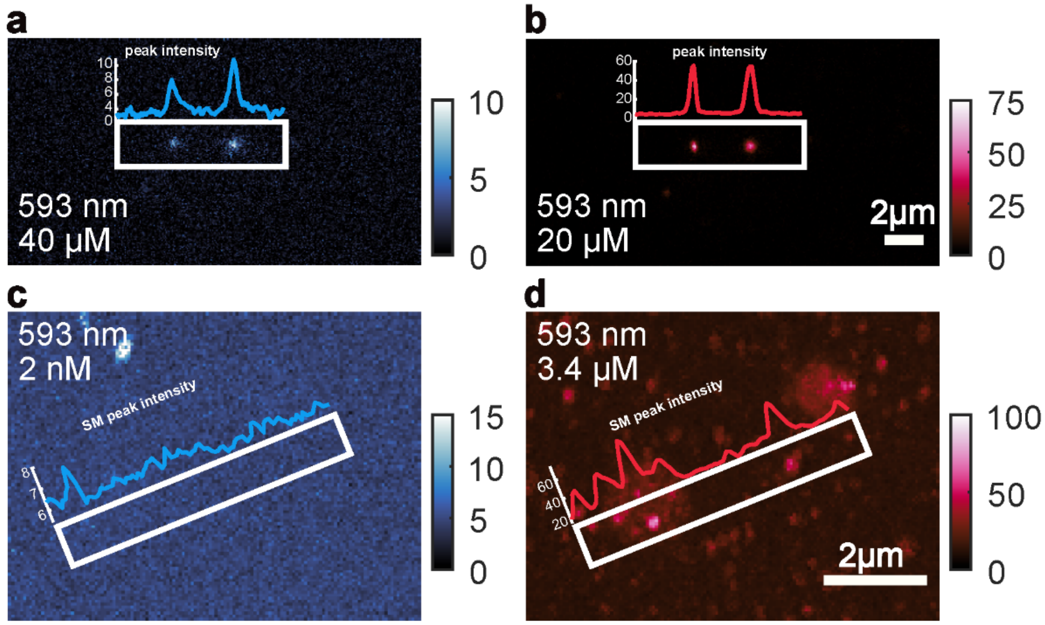

**Fig. S5. Sequential Nile blue-Nile red (NB-NR) imaging at epifluorescence and single-molecule concentrations within the orange (593 nm) emission window. a,b**: Imaging (a) 40 µM NB followed by (b) 20 µM NR within the same condensates using identical 532-nm excitation and 20 ms exposure time. **c,d**: Maximum intensity projections of 2000 frames of (c) 2 nM NB followed by (b) 3.4 µM NR within the same condensates using identical 562-nm excitation and 10 ms exposure time. Note that pumping NB at these wavelengths is far off resonance, resulting in very weak NB fluorescence, which is ~12-fold weaker than NR at the same epifluorescence concentration. At single-molecule concentrations, individual NB "spots" are not detectable, while many NR bright "spots" are seen within these condensates.

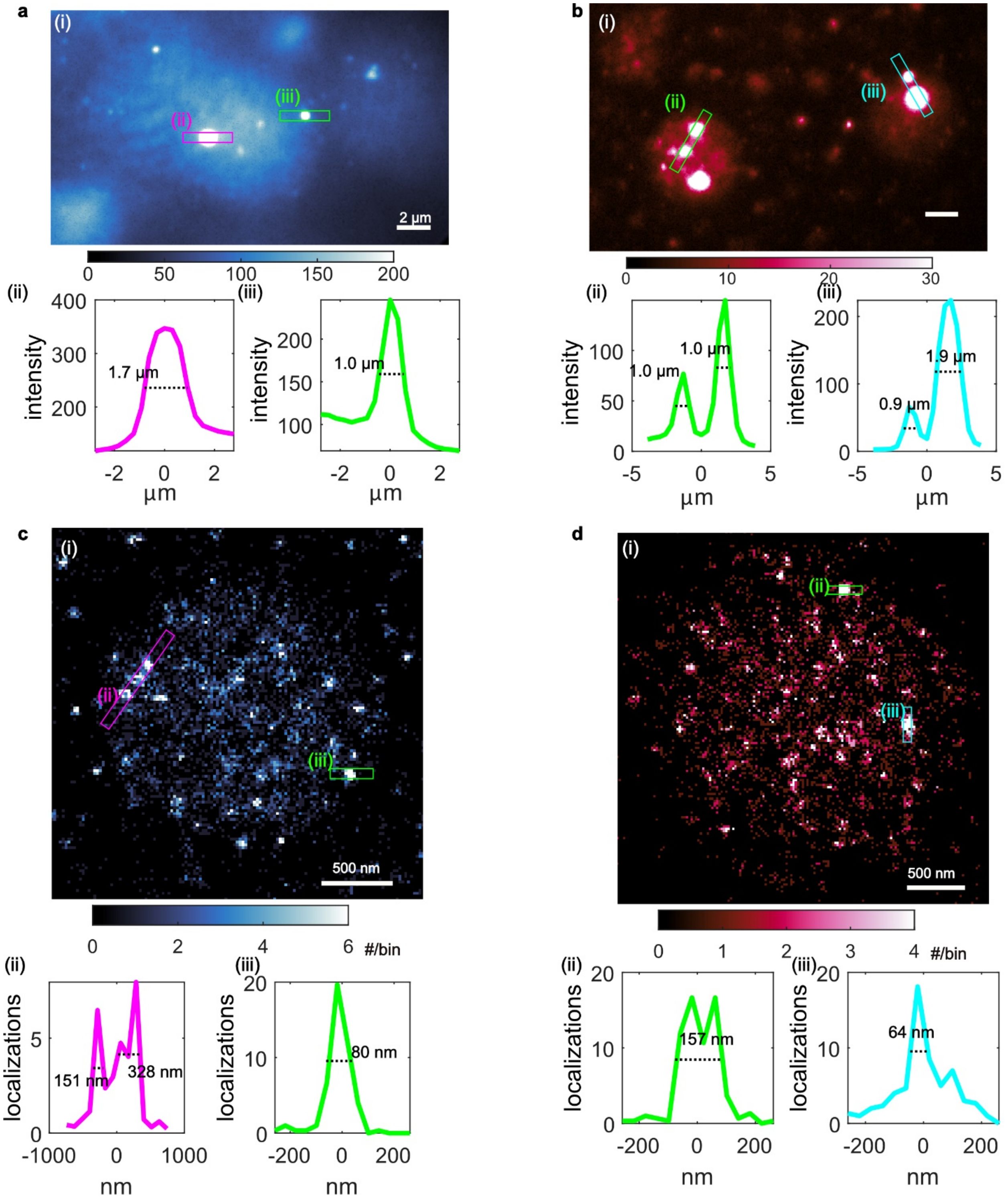

**Fig. S6. Microscale and nanoscale hubs measured by epifluorescence imaging and SMLM. a**, **b**: epifluorescence images of A1-LCD condensates using (**a**) NB and (**b**) NR, reproduced from **Fig. 1(b)**. **c**, **d**: SMLM image of A1-LCD condensates using (**c**) NB and (**d**) NR, reproduced from **Fig. 3g** and **Extended Data Fig. 5g**. (ii, iii): intensity or localization profile along the long axes of boxes shown in (i) labelled with full-width at half-maximum.

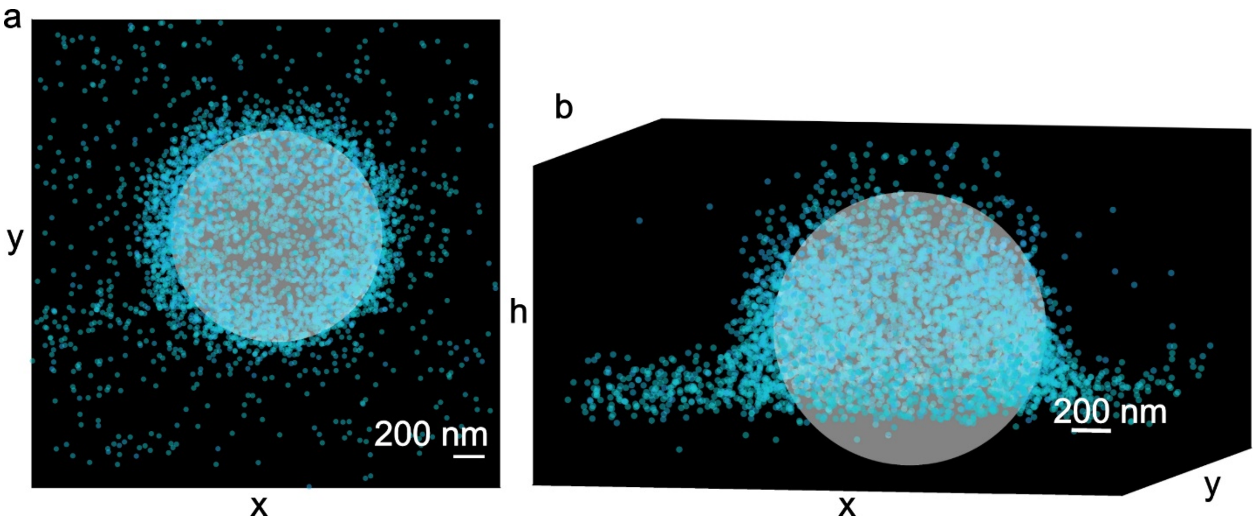

**Fig. S7. Fitting the 3D condensate structure with an ellipsoid. a**: The x-y view and (**b**) 3D view of the 3D position of MC540 and fitted ellipsoid.

**Supplementary Movies**

MovieS1_SM_reconstruction.mp4

**Movie S1. Detecting and localizing single NR molecules within an A1-LCD condensate.** Left: Position estimates (green crosses) overlaid on the raw SM images. Right: SMLM reconstruction formed by accumulating localizations over time. Each detected single molecule is represented as a shaded red disk.

MovieS2_NR_sequence_video_labelled.mp4

**Movie S2. SMLM images of NR within an A1-LCD condensate reconstructed using subsets of raw SM images**. The hubs of NR disappear and appear at nearby positions over time (sec). Yellow circles: hubs

MovieS3_NB_sequence_video_labelled.mp4

**Movie S3. SMLM images of NB within an A1-LCD condensate reconstructed using subsets of raw SM images**. The hubs of NB disappear and appear at nearby positions over time (sec). Yellow circles: hubs.

MovieS4_SM_tracking.mp4

**Movie S4. Single-molecule tracking of NB within A1-LCD condensates**. Yellow crosses represent the position of each SM, while yellow lines represent the trajectory of each SM overlaid on the raw SM images.

MovieS5_MC540_orientation.mp4

**Movie S5. Single-molecule orientation-localization microscopy of MC540 localized at the interfaces of LCD condensates**: (left) $Aro^-$, (middle) A1-LCD (WT), and (right) $Aro^+$ variants. Each molecule is depicted as a line segment whose center represents the xy position of MC540 and whose long axis points along the orientation of MC540's emission dipole measured by the pixOL microscope. Several slices along z are shown for MC540 located at different heights h above the coverslip.
